# Chemical foundation models reveal molecular principles underlying glycobiology

**DOI:** 10.64898/2026.10.13.758502

**Authors:** Manuel Sokolov Ravasqueira, João Bettencourt-Silva, Vanessa López García, Raúl Fernández-Díaz, Natasha Mulligan, Gavin P. Davey

**Author notes:** Correspondence should be sent to: Manuel Sokolov Ravasqueira and Gavin Davey.

## Abstract

Glycans encode biological specificity through branching, linkage and stereochemical diversity, and have traditionally been modelled using specialised computational representations distinct from those developed for general chemistry. It remains unclear to what extent representations learned from general molecular chemistry capture the molecular principles of glycobiology. Here, we systematically evaluate the transfer of chemical foundation models to glycobiology using standardized benchmarks spanning taxonomy, immunogenicity, glycosylation type and protein-glycan interactions. We show that atom-level representations learned from general chemistry encode biologically meaningful molecular organization, providing particularly strong performance for molecular recognition, including immunogenicity and protein-glycan interaction prediction. Fine-tuning reorganizes these representations into latent spaces aligned with glycan architecture, glycosylation programmes and molecular recognition. Mechanistic analyses identify stereochemistry as an important determinant of successful transfer, while tokenization, context length and conformer generation influence the representation of large, branched glycans. These representations generalize from natural glycans to synthetic glycomimetics and recover experimentally validated stereochemistry-dependent glycan-lectin recognition. Together, our findings demonstrate that molecular principles learned from general chemistry are sufficient to capture fundamental features of glycobiology. They position glycans as a specialised region of chemical space and chemical foundation models as a broadly applicable framework for glycan function prediction, molecular recognition and glycomimetic discovery.

## Introduction

Glycans are among the most structurally diverse biomolecules, regulating protein folding, immune recognition, cell-cell communication and host-pathogen interactions through highly branched carbohydrate structures^1^. Unlike proteins and nucleic acids, glycans are assembled by coordinated networks of glycosyltransferases and glycosidases rather than template-directed biosynthesis, generating extensive branching, linkage diversity and stereochemical complexity^2,3^. Glycans exist as free oligosaccharides or are covalently attached to proteins, lipids, RNAs, and other glycans, where they influence molecular stability, trafficking and biological recognition ^2,4^.

Glycan biosynthesis is tightly integrated with cellular metabolism, relying on nucleotide-sugar donors generated by central metabolic pathways and responding dynamically to nutrient availability, energy status and metabolic flux, thereby linking cellular metabolism to molecular recognition and signalling ^5–8^. More than 60% of human proteins are glycosylated, while altered glycosylation is a hallmark of cancer, inflammatory disorders and infectious disease ^9–11^. Consequently, advances in glycomics technologies and curated glycan databases have created growing opportunities for applying artificial intelligence and in silico methods in glycobiology ^12–24^.

The structural complexity of glycans has led them to be treated as a distinct computational problem. Glycoinformatics has consequently relied on carbohydrate-specific representations, including IUPAC-Condensed sequences, WURCS, GlycoCT and graph-based encodings that explicitly preserve monosaccharide identities, glycosidic linkages and branching patterns ^25,26^. More recently, glycans have also been represented at atomic-level using SMILES. These representations have supported the development of numerous glycan-specific machine-learning models ^27–33^, and the GlycanML benchmark, which evaluates taxonomy, immunogenicity, glycosylation-type and protein-glycan interaction prediction ^34^.

In parallel, chemical foundation models (ChemFMs) have expanded molecular machine learning by learning transferable atom-level representations from large collections of chemical structures. Transformer-based ChemFM models including ChemBERTa, MoLFormer, and MAMMAL together with 3-dimensional models such as Uni-Mol learn transferable atom-level representations from SMILES strings or molecular conformations through large-scale self-supervised pretraining ^35–38^. These models have achieved strong performance across diverse molecular-prediction tasks and can transfer chemical knowledge between related domains^39–42^. However, glycans remain underrepresented in general chemical data resources^43^, leaving unresolved whether chemical foundation models are exposed to sufficient glycan diversity during pre-training and whether the resulting representations retain the structural information required to model glycan biology.

Despite their biological complexity, glycans remain chemical molecules governed by the same physicochemical principles as other biomolecules. Molecular recognition depends on atom-level chemistry, stereochemistry and local molecular environments, the same features explicitly represented by modern chemical foundation models. Unlike glycan-specific representations, atom-level molecular representations also naturally accommodate synthetic substitutions and non-natural modifications found in glycomimetics. This raises a central question: can models trained on general chemical space capture the molecular principles governing glycobiology, or do glycans require specialised representations?

Resolving this question requires more than comparing predictive performance. Evidence of successful transfer should establish that chemical foundation models retain glycan-defining information, including stereochemistry, linkage, branching and molecular context; that biological supervision produces representations aligned with glycan structure and function; and that the resulting models generalize beyond the natural glycans used for training. These criteria distinguish genuine transfer of chemically meaningful information from task-specific predictive performance.

Here, we systematically evaluate the transfer of chemical foundation models to glycobiology using benchmarks spanning taxonomy, immunogenicity, glycosylation type and protein-glycan interactions. Combining comparative benchmarking, mechanistic analyses of representation learning and independent experimental validation, we show that general chemical foundation models recover transferable molecular principles underlying glycobiology. Fine-tuning reorganizes pretrained chemical representations into biologically coherent latent spaces, stereochemistry emerges as the principal determinant of successful transfer, and learned representations generalize from natural glycans to synthetic glycomimetics while recovering experimentally validated glycan-lectin recognition.

## Results

### General chemical representations transfer effectively to glycobiology

We evaluated the ability of atom-level molecular representations learned from general chemistry to transfer to glycan biology under two complementary regimes, static representations and task-adaptive representations (Figure 1a). Static representations comprised fixed engineered descriptors, molecular fingerprints^44^, and embeddings extracted from pretrained chemical foundation models, whereas task-adapted representations additionally included chemical foundation models fine-tuned for each downstream task together with specialist glycan models trained directly on glycan-native sequence or graph representations. All models were evaluated using the GlycanMLBenchmark ^34^ across glycan taxonomy, immunogenicity, glycosylation-type prediction, and protein-glycan interaction prediction (*see Methods).* To enable direct comparison between glycan-native and general purpose chemical representations, glycans were converted from IUPAC-condensed notation to SMILES using GlyLES ^45^ while preserving the benchmark training, validation and test partitions: motif-based splits for the glycan-only tasks and protein-sequence-similarity-based splits for protein-glycan interaction prediction (Supplementary Table 1 and Supplementary Figs. 1 and 2).

**Figure 1.**
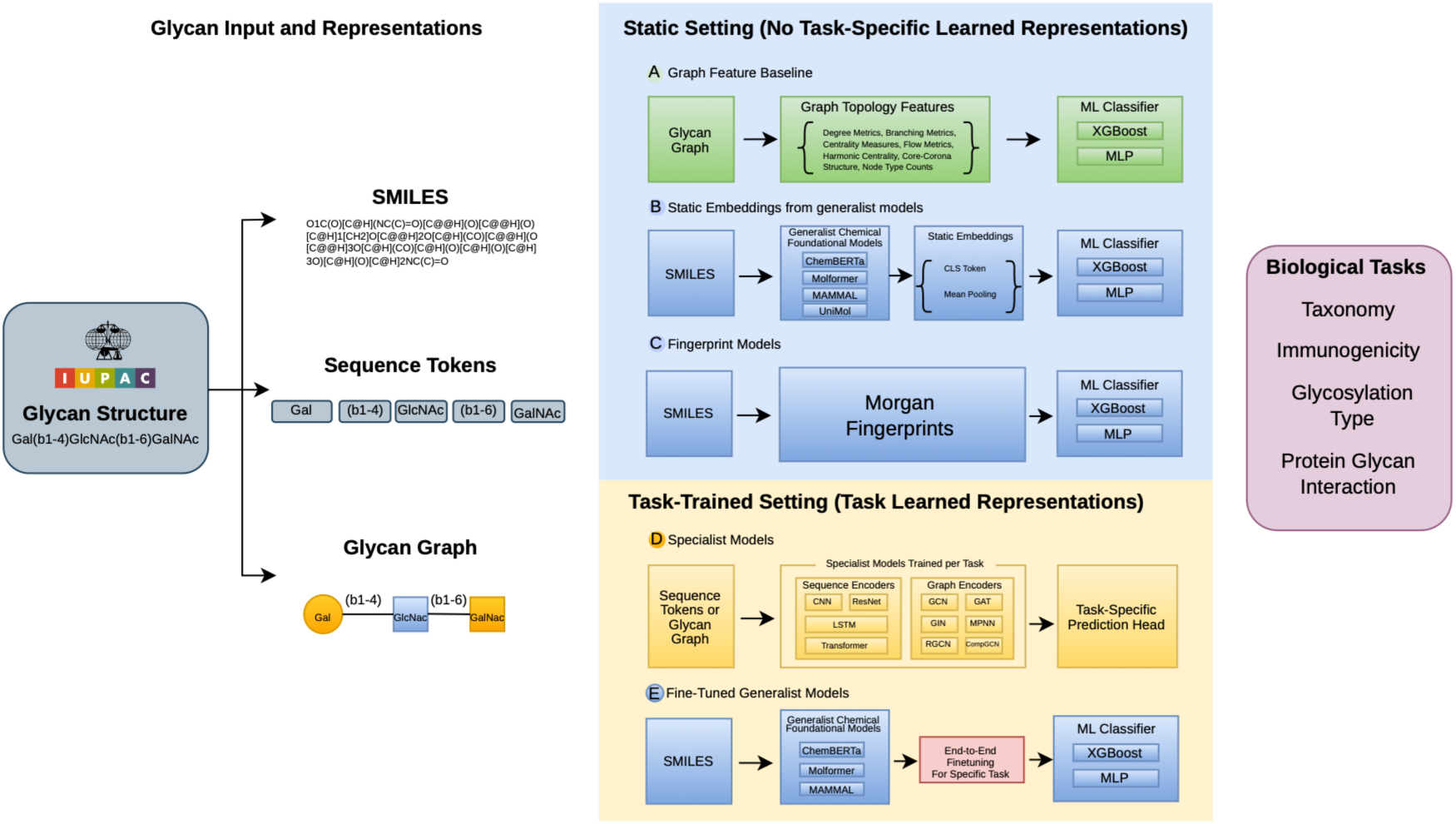
Benchmarking framework for evaluating chemical foundation model transfer to glycobiology. Glycans were represented as molecular SMILES, glycan sequences, or glycan graphs and evaluated using two complementary modelling regimes. In the static setting, fixed representations, including (A) handcrafted graph descriptors, (B) frozen embeddings from chemical foundation models (ChemBERTa, MoLFormer, MAMMAL and Uni-Mol) and (C) Morgan fingerprints, were combined with downstream XGBoost or MLP classifiers. In the task-trained setting, (D) specialist glycan models trained on sequence or graph representations and fine-tuned chemical foundation models were optimized end-to-end for each prediction task. Blue pathways indicate SMILES-based atomic-level molecular representations. Models were benchmarked across glycan taxonomy, immunogenicity, glycosylation-type and protein-glycan interaction prediction.

We first examined the static setting, where molecular representations remained fixed and only downstream prediction models XGBoost or MLP were trained (Supplementary Table 2). Morgan fingerprints were particularly effective for taxonomic prediction. Their advantage was most apparent for glycan taxonomy, where ECFP4 and ECFP6 fingerprints, combined with XGBoost or MLP classifiers, achieved the highest performance for both high-level (macro-F1 = 0.625) and low-level taxonomy (macro-F1 = 0.343). These results are particularly notable because the original taxonomy benchmark deliberately introduces substantial train-test shifts in glycan size and structural complexity (Fig. 2a,b), requiring representations to generalise beyond closely related structures rather than memorising similar glycans. The strong performance of Morgan fingerprints therefore demonstrates that much of the information required for glycan structural classification is already captured by local atom-centred molecular environments.

**Figure 2.**
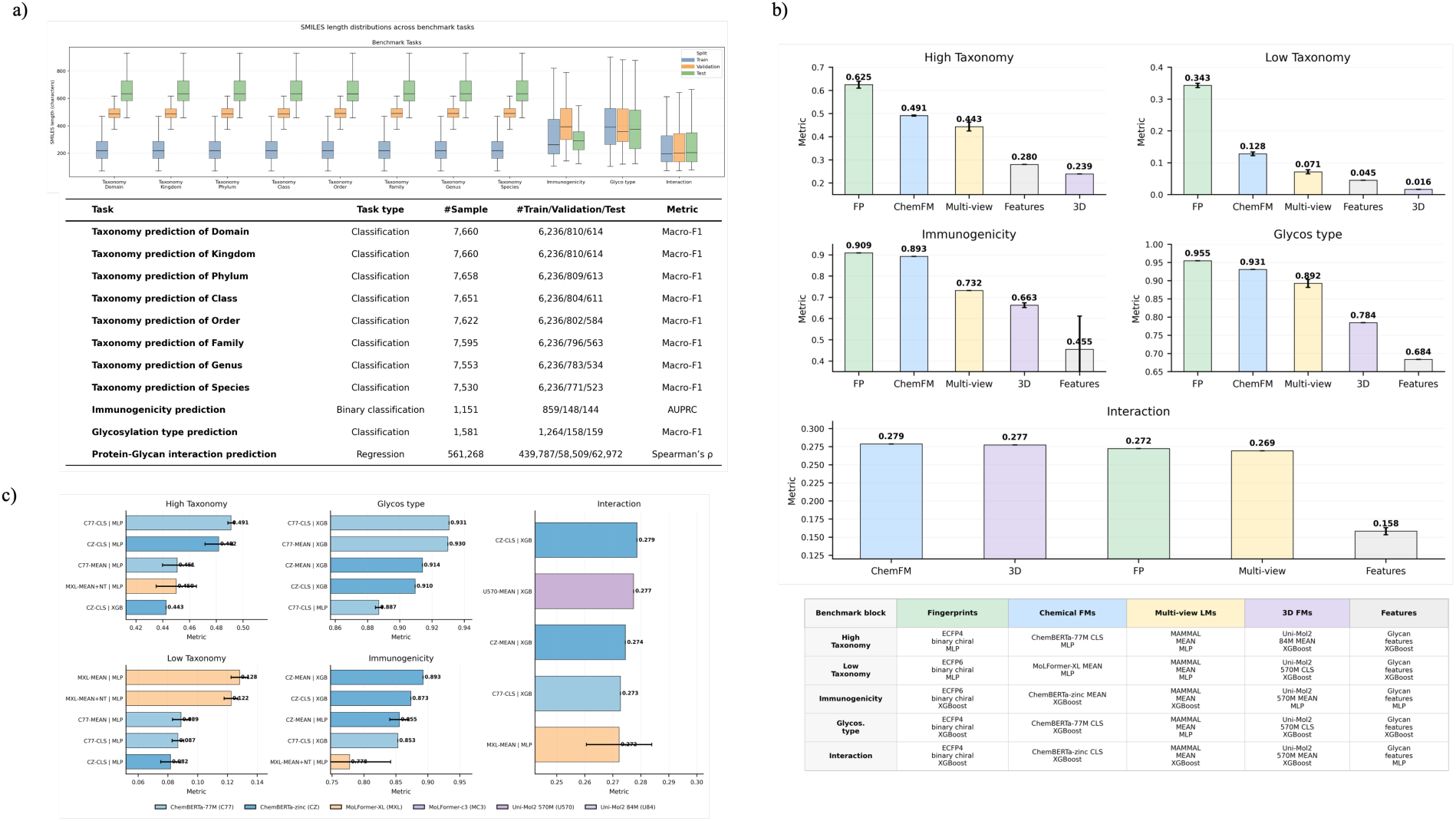
Static benchmarking reveals complementary strengths of molecular representations for glycan prediction. **a)** Dataset overview showing SMILES length distributions, task types, sample sizes, train/validation/test splits, and evaluation metrics. **b)** Aggregated comparison by best of each representation family, showing that Morgan fingerprints dominate taxonomic prediction and remain strong for immunogenicity and glycosylation type, whereas chemical and 3D foundation model embeddings are most competitive for protein-glycan interaction prediction. **(c)** Best frozen chemical foundation model configurations per task, highlighting their competitive functional performance and strongest static interaction signal. Error bars indicate variability across repeated evaluations or splits where applicable.

Frozen chemical foundation model embeddings provided the strongest alternative to fingerprints in the static setting. Among the evaluated models, ChemBERTa-family encoders consistently produced the strongest transferable molecular representations (Figure 2c). Although they did not surpass Morgan fingerprints for taxonomy, ChemBERTa embeddings substantially outperformed handcrafted glycan graph descriptors and other frozen molecular foundation models while approaching fingerprint performance for both immunogenicity (AUPRC = 0.893) and glycosylation-type prediction (macro-F1 = 0.931). These findings indicate that molecular representations learned entirely from conventional chemical corpora retain biologically meaningful information relevant to glycans despite the absence of glycan-specific pretraining.

Protein-glycan interaction prediction presented a more stringent test because the benchmark provides a protein-similarity based split across training and test sets (Supplementary Fig. 2). For example, H1N1 haemagglutinins interactions are included during training, whereas H5N5 haemagglutinin interactions are held out for testing, requiring transfer across influenza subtypes. Despite the modest correlations across all models, frozen ChemBERTa-zinc CLS embeddings combined with XGBoost achieved the highest static performance (Spearman’s ρ = 0.279), outperforming Morgan fingerprints and handcrafted glycan descriptors (Supplementary Table 2, Fig. 2c) with ChemBERTa providing the strongest transferable static representation.

Additionally, systematic comparison of modelling choices revealed that CLS pooling significantly improved low-level taxonomy across chemical foundation models, ECFP6 fingerprints consistently outperformed ECFP4, and downstream classifier performance was task dependent, with MLP favouring taxonomy whereas XGBoost achieved superior performance for immunogenicity, glycosylation-type and protein-glycan interaction prediction (Supplementary Fig. 3).

Together, the static benchmark establishes two key observations. First, Morgan fingerprints provide an exceptionally strong baseline for glycan structural classification. Second, ChemBERTa-family encoders transfer substantial molecular knowledge to glycans, approaching fingerprint performance for functional prediction, while providing the strongest frozen representations for protein-glycan interactions. Unlike fingerprints, a general purpose static representation, chemical foundation models can be refined through end-to-end learning, prompting us to next examine whether task-specific fine-tuning further reorganises molecular representations to capture glycan function.

### Task adaptation reveals a structural-functional divide in glycan representation learning

Having established the performance of fixed molecular representations, we next asked whether task-specific representation learning could further improve glycan prediction. We compared fixed representations with two trainable representation-learning strategies: specialist glycan models trained on glycan-native IUPAC-condensed (monomer and glycosidic linkage representations) sequences or glycan graphs, and fine-tuned chemical foundation models in which the molecular encoder was adapted for each downstream task (Figure 3).

**Figure 3.**
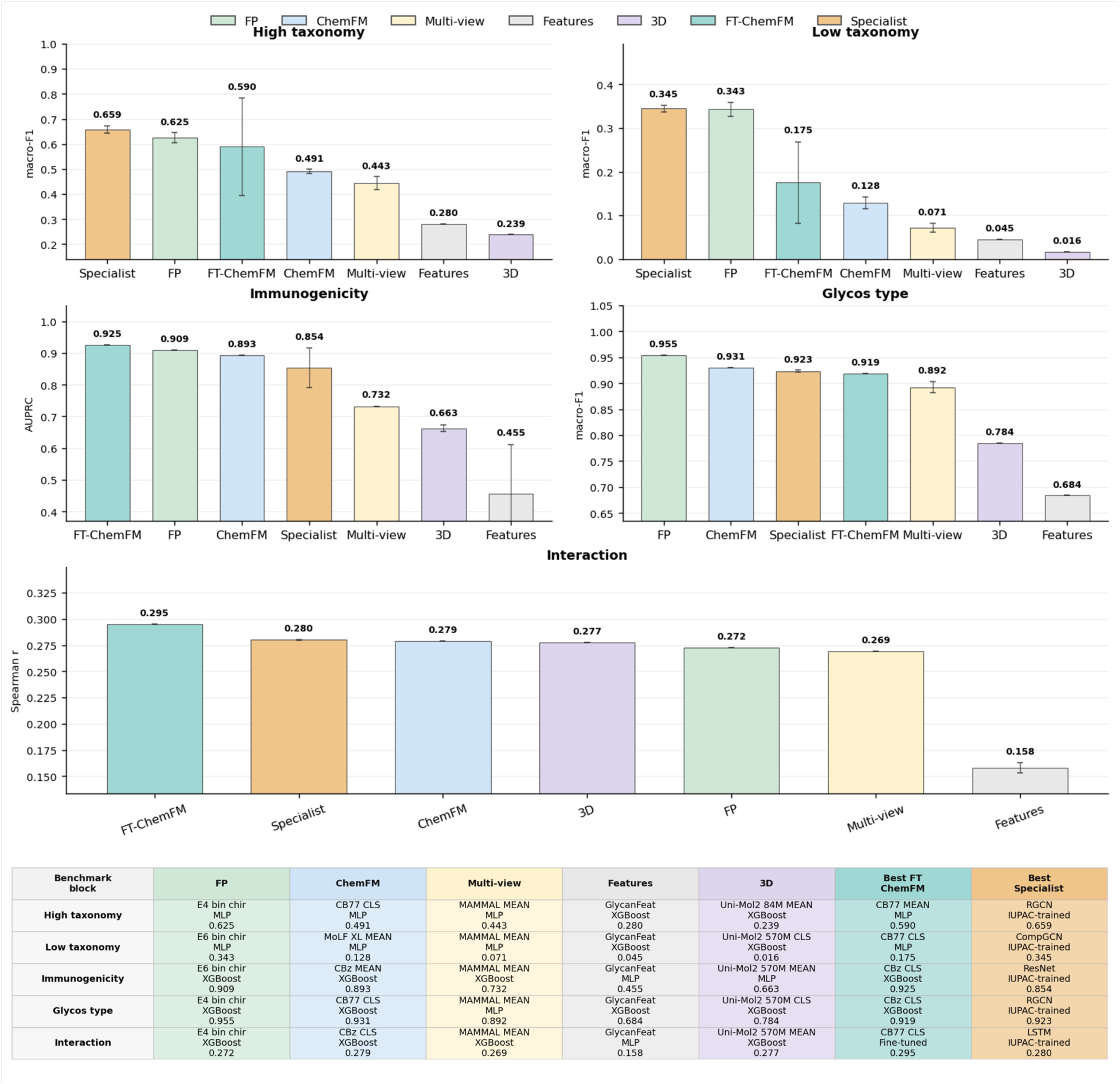
Task adaptation reveals a structural-functional divide in glycan representation learning. For each benchmark task, the best model from each representation family is shown, including Morgan fingerprints (FP), frozen ChemFM embeddings, multi-view molecular models, handcrafted glycan graph features, 3D molecular foundation models, fine-tuned chemical foundation models (FT-ChemFM), and specialist glycan models trained on glycan-native sequence or graph representations. Bars report the best observed performance per family for high-level taxonomy, low-level taxonomy, immunogenicity, glycosylation type, and protein-glycan interaction prediction, using macro-F1, AUPRC, or Spearman’s correlation as appropriate. The accompanying table summarizes the best-performing model configuration for each representation family and task. Error bars indicate variability across repeated evaluations.

A clear distinction emerged between structural and functional prediction tasks. Glycan taxonomy was best captured by specialist glycan models trained on IUPAC-condensed sequences or glycan graphs, with the strongest model achieving macro-F1 values of 0.659 for high-level and 0.345 for low-level taxonomy. This likely reflects the explicit preservation of monosaccharide identity and linkage information within glycan-native representations. Nevertheless, the static Morgan fingerprints remained highly competitive (macro-F1 = 0.625 and 0.343, respectively), indicating that local atom-centred molecular environments capture much of the information required for glycan structural classification.

Fine-tuning also substantially improved chemical foundation model performance, particularly for high-level taxonomy, where the best ChemFM increased from a macro-F1 of 0.491 to 0.590. Low-level taxonomy also improved (0.128 to 0.175), indicating that biological supervision better aligns pretrained molecular representations with glycan structural organization and narrows the gap with specialist glycan models. These improvements were model dependent: smaller ChemBERTa-family encoders benefited most from fine-tuning, whereas larger models such as MoLFormer generally showed reduced gains (Supplementary Figs. 4 and 5).

For functional prediction tasks, the pattern reversed in favour of general-purpose molecular representations. Fine-tuned ChemBERTa foundation models achieved the best immunogenicity performance (AUPRC = 0.925), outperforming Morgan fingerprints (0.909) and specialist glycan models (0.854). They also achieved the strongest protein-glycan interaction prediction, (Spearman’s correlation = 0.295), exceeding specialist glycan models (0.280), frozen ChemFM embeddings (0.279), 3D embeddings (0.277), and fingerprints (0.272). These findings indicate that atom-level molecular representations become increasingly advantageous when prediction depends upon molecular recognition rather than glycan structural identity.

Glycosylation-type prediction exhibited an intermediate pattern. Morgan fingerprints remained the strongest representation (macro-F1=0.955), while frozen ChemBERTa embeddings (0.931) slightly outperformed specialist glycan models (0.923), suggesting that this task is largely determined by local molecular substructures and requires less task-specific adaptation.

Collectively, these results reveal a clear structural-functional divide in glycan representation learning. Specialist glycan models remain strongest for taxonomy, whereas ChemBERTa-family models, particularly after fine-tuning, provide the best representations for functional prediction and protein-glycan interactions. These findings demonstrate that molecular knowledge learned during chemical pretraining can be successfully adapted to capture glycan-specific biological function. Full Results and Parameters including static and task-adapted setting provided in Supplementary Tables 3 and 4.

### Fine-tuning reorganizes transferable chemical representations into biological programmes

Having established that ChemBERTa provides transferable representations for glycans, we next asked whether biological supervision simply improves prediction accuracy or fundamentally reorganizes the underlying chemical representation space. We compared pretrained and fine-tuned ChemBERTa embeddings using Leiden clustering in the original high-dimensional space (Fig. 4). Clusters were identified independently of biological labels and subsequently characterized by benchmark annotations and cluster-versus-rest motif enrichment (Supplementary Fig. 6) and, for protein-glycan interactions, glycan embedding cluster-protein binding enrichment. Across all benchmark tasks, pretrained embeddings already exhibited biologically meaningful chemical organization, whereas fine-tuning consistently refined these broad chemical neighbourhoods into communities more closely aligned with the corresponding biological functions.

**Figure 4.**
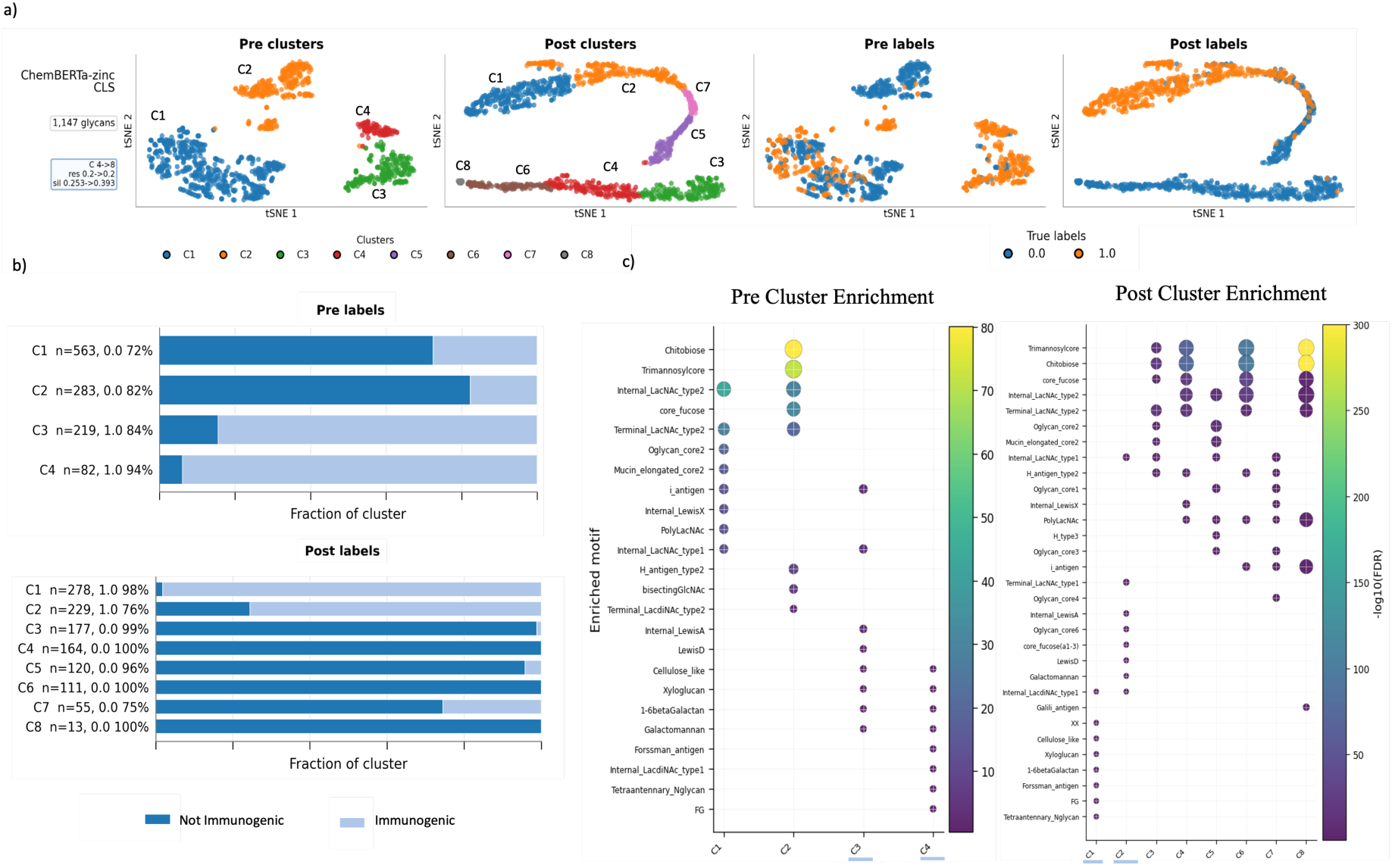
Fine-tuning reorganizes chemical representations into immunogenicity-associated glycan communities. **a)** t-SNE projections of ChemBERTa-zinc CLS glycan embeddings before and after immunogenicity fine-tuning. Leiden communities were identified in the original high-dimensional representation space and projected onto t-SNE for visualization. Plots are coloured either by Leiden community or by the benchmark immunogenicity label (0, non-immunogenic; 1, immunogenic). **b)** Immunogenicity-label composition of each Leiden community. Bars show the fraction of immunogenic and non-immunogenic glycans, with community size and the dominant-label. **c)** Cluster-versus-rest enrichment of annotated glycan motifs before and after fine-tuning. Bubble plots show significantly enriched glycan motifs within each Leiden community; colour indicates statistical significance (-log10(FDR)), and bubble size represents enrichment magnitude. Fine-tuning reorganized broad chemical communities into more distinct immunogenic and non-immunogenic structural programmes. Underscores beneath the cluster labels indicate communities classified as immunogenic based on their dominant label.

#### Fine-tuning sharpens latent self/non-self organization

Immunogenicity provided the clearest example of task-driven representation reorganization (Fig. 4). Before fine-tuning, ChemBERTa-zinc embeddings separated glycans into four communities that broadly distinguished mammalian-associated glycans from plant-, microbial- and rare-sugar-rich structures (Fig. 4; Supplementary Fig. 7). The two largest non-immunogenic communities (C1, 72% and C2, 82% non-immunogenic) were enriched for canonical N-glycan, LacNAc and mucin-associated architectures, whereas the two immunogenic communities (C3, 84% and C4, 94% immunogenic) captured plant-, microbial- and xeno-antigen-associated carbohydrate programmes together with rare-sugar motifs. Thus, pretrained embeddings already encoded a broad self/non-self organization despite the absence of glycan-specific training.

Fine-tuning preserved this global organization while substantially increasing its biological resolution. Immunogenic glycans became concentrated within two communities (C1, 98% and C2, 76% immunogenic), whereas the broader non-immunogenic space resolved into multiple canonical N-glycan, LacNAc/polyLacNAc and mucin-type O-glycan programmes. Rather than creating new structural relationships, fine-tuning refined existing chemical neighbourhoods into more coherent biological communities, increasing both label separation and structural resolution while maintaining the underlying distinction between mammalian and foreign-associated glycans (Fig. 4; Supplementary Fig. 7).

#### Glycosylation type emerges as distinct molecular grammar

Before fine-tuning, ChemBERTa-zinc embeddings already captured the fundamental distinction between N- and O-glycan biosynthetic grammar (Supplementary Figs. 8 and 9). Two broad communities predominated: an O-linked community enriched for mucin-type glycans and a predominantly N-linked community enriched for canonical N-glycan architecture. Thus, pretrained embeddings already separated GalNAc-centred O-glycan chemistry from Man/GlcNAc-centred N-glycan chemistry.

Fine-tuning preserved this global organization while substantially increasing its structural resolution. N-linked glycans separated into distinct canonical, branched and extended N-glycan communities (99-100% N-linked), whereas O-linked glycans resolved into core-1, core- 2/extended and terminally elaborated mucin-type programmes (99-100% O-linked). Free glycans formed a dedicated community, while a single mixed community retained terminal LacNAc- and sialylation-associated structures shared across attachment classes. Thus, fine-tuning refined broad N- and O-glycan programmes into discrete biosynthetic communities while preserving their underlying molecular relationships (Supplementary Figs. 8 and 9).

#### Broad chemical chemotypes reorganize into kingdom-associated glycan ecologies

A similar pattern emerged for kingdom classification. Before fine-tuning, ChemBERTa-77M embeddings showed only coarse alignment with taxonomy, separating common eukaryotic glycan scaffolds from a chemically distinct non-mammalian carbohydrate space enriched for plant-, bacterial- and other non-vertebrate glycans (Supplementary Figs. 10 and 11). Thus, pretrained representations primarily captured broad molecular chemotypes rather than kingdom-specific organization.

Fine-tuning preserved these underlying chemical relationships while reorganizing them into more taxonomically coherent communities. Distinct bacterial (94%) and animal (95%) communities emerged, characterized by bacterial cell-envelope glycans and canonical animal N- and O-glycan architectures, respectively, whereas plant glycans formed a separate polysaccharide-associated community. Fungal glycans remained a more heterogeneous intermediate group, consistent with their structural overlap with plant and other non-vertebrate carbohydrate architectures. Thus, biological supervision refined broad chemical neighbourhoods into biologically meaningful kingdom-associated glycan ecologies without disrupting their underlying molecular organization (Supplementary Figs. 10 and 11).

#### Protein-glycan interaction fine-tuning reveals molecular recognition programmes

Protein-glycan interaction prediction required a modified analysis because benchmark labels are defined for glycan-protein pairs rather than individual glycans. Glycans were assigned to Leiden communities before and after fine-tuning, and protein-specific interaction enrichment was assessed for each community to identify glycan structural programmes preferentially recognized by different proteins. Before fine-tuning, ChemBERTa-77M embeddings separated glycans into six chemically defined communities that already exhibited biologically plausible protein associations (Supplementary Figs. 12 and 13). Notably, influenza haemagglutinins preferentially associated with sialylated glycan communities, consistent with their established recognition of sialic-acid-containing receptors. Fine-tuning preserved these pre-existing relationships while reorganizing them into three more coherent recognition programmes corresponding to terminal fucosylated-Lewis-associated glycans, sialylated Gal-LacNAc-rich receptor glycans, and core N-glycan-scaffold structures. These programmes were associated with distinct groups of lectins, microbial adhesins and viral glycoproteins, producing more specific and biologically interpretable protein-cluster relationships (Supplementary Figs. 12 and 13).

Thus, task supervision refined rather than created molecular recognition patterns, concentrating broadly distributed chemical features into coherent glycan recognition programmes while preserving established interactions such as the association between influenza haemagglutinins and sialylated glycans.

Collectively, these analyses reveal a common principle: pretrained ChemBERTa embeddings already encode substantial glycobiologically relevant chemical organization, and fine-tuning consistently refines rather than replaces these relationships. Across all benchmark tasks, broad chemical neighbourhoods reorganize into biologically coherent communities corresponding to immunogenicity, glycosylation type, kingdom and protein-recognition programmes. This representation-level reorganization provides a mechanistic explanation for improved prediction, demonstrating that biological supervision concentrates task-relevant structural information into more interpretable molecular programmes. Complete cluster compositions and enrichment analyses are provided in Supplementary Table 5.

### Stereochemistry governs transferable glycan representations

The preceding analyses demonstrated that fine-tuning selectively improves functional prediction and reorganizes molecular representation space into biologically coherent glycan communities. We next asked which molecular features within the atom-level representation contribute to these improvements. Because glycan recognition is fundamentally dependent on stereochemistry, we performed a stereochemistry ablation in the static setting by removing stereochemical annotations glycan SMILES strings before embedding extraction from chemical foundation models, while recomputing Morgan fingerprints without chirality information. Downstream classifiers, benchmark splits, and evaluation metrics were kept identical, enabling paired comparison between chiral and achiral molecular representations (Fig. 5).

**Figure 5.**
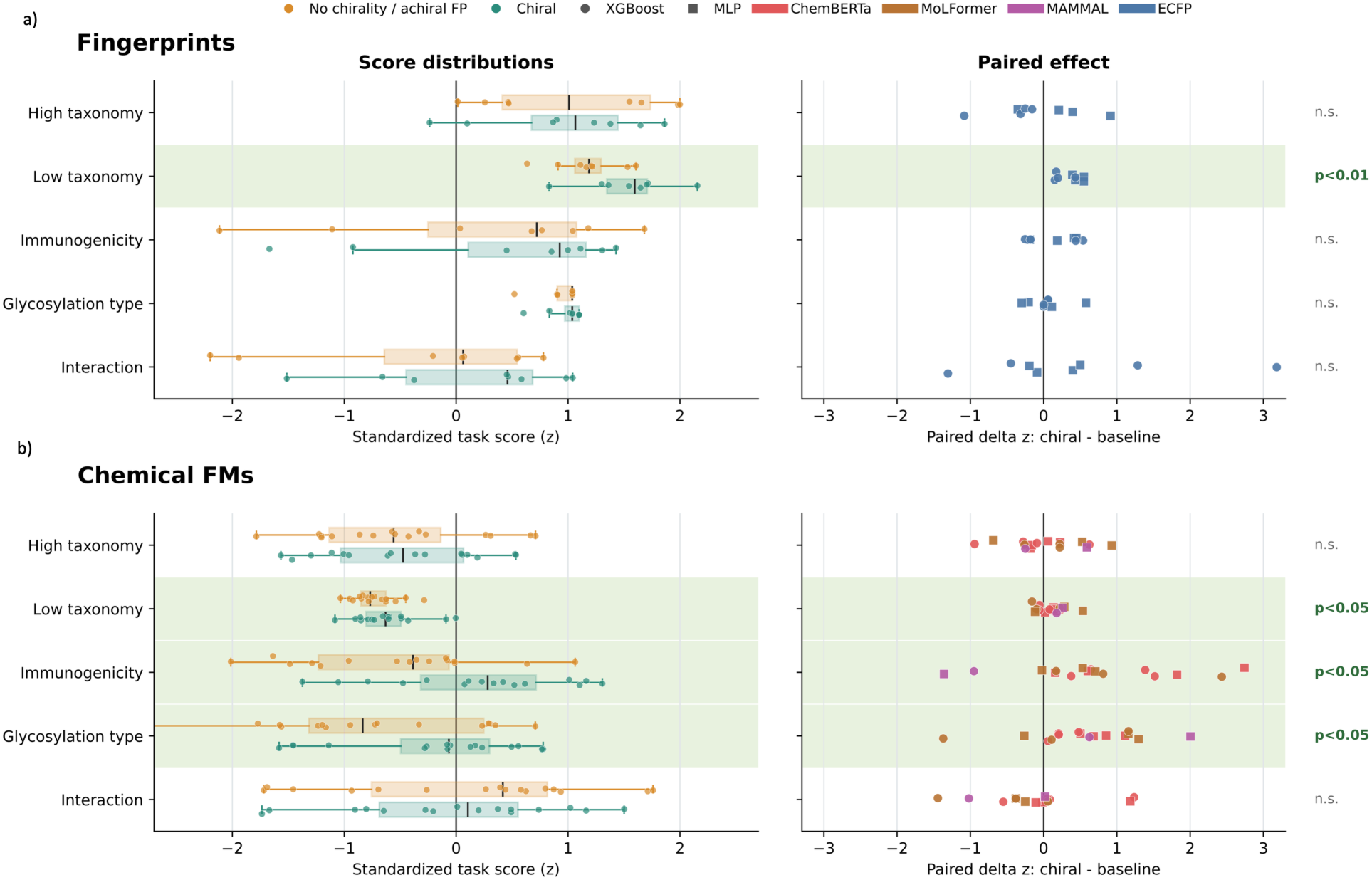
Stereochemistry is an important determinant of transferable glycan representations. **(a)** Effect of removing chirality from Morgan fingerprints across benchmark tasks. Left, standardized task-score distributions for chirality-aware and achiral fingerprints across downstream classifiers. Right, paired performance differences (chiral-achiral) for each model/classifier combination. Chirality significantly improved low-level taxonomy prediction but showed no significant effect for the remaining benchmark tasks. **(b)** Effect of removing stereochemical annotations from SMILES before extracting chemical foundation model embeddings. Left, standardized performance distributions for chiral and achiral SMILES representations. Right, paired performance differences for each model/classifier combination. Stereochemistry significantly improved low-level taxonomy, immunogenicity and glycosylation-type prediction, whereas high-level taxonomy and protein-glycan interaction prediction showed no significant effect. Points represent individual model/classifier combinations; colours denote foundation model families and symbols denote downstream classifiers. Shaded regions indicate statistically significant paired effects.

Chemical foundation models were significantly affected by the presence of stereochemical information. Across model/head combinations, chiral SMILES representations produced consistent performance gains over achiral SMILES for low-level taxonomy, immunogenicity, and glycosylation-type prediction. Importantly, including chirality did not significantly reduce performance for any benchmark task; rather, when statistically significant paired effects were observed, they consistently favoured stereochemistry-aware representations. These findings demonstrate that atom-level molecular representations capture stereochemical information that contributes predictive signal beyond molecular composition and connectivity alone.

This effect was model-dependent. For ChemBERTa-family models, the improvement with chiral SMILES suggests that stereochemical tokens contribute directly to the learned molecular representation and improve downstream glycan prediction. By contrast, MoLFormer was pretrained primarily on canonicalized SMILES that lack explicit stereochemical annotations.

Consequently, changes in MoLFormer performance following stereochemistry ablation likely reflect altered input representations rather than stereochemical knowledge acquired during pretraining. Collectively, these results demonstrate that successful transfer from general chemical foundation models to glycobiology depends not only on atom-level molecular chemistry, but also on preserving the stereochemical information that distinguishes biologically distinct carbohydrate structures.

### Architectural determinants of successful transfer to glycobiology

To understand why current molecular foundation models transfer unevenly to glycans, we examined how glycan SMILES are processed before representation learning. We focused on model-specific tokenization, sequence truncation, chirality handling, and 3D conformer generation (Fig. 6). Because truncation occurs after tokenization, identical glycans may be fully retained by one model but partially discarded by another depending on tokenizer vocabulary and maximum input length. This is particularly important for glycans, which are typically long, branched and stereochemically rich compared with the small molecules used to pretrain most chemical foundation models.

**Figure 6.**
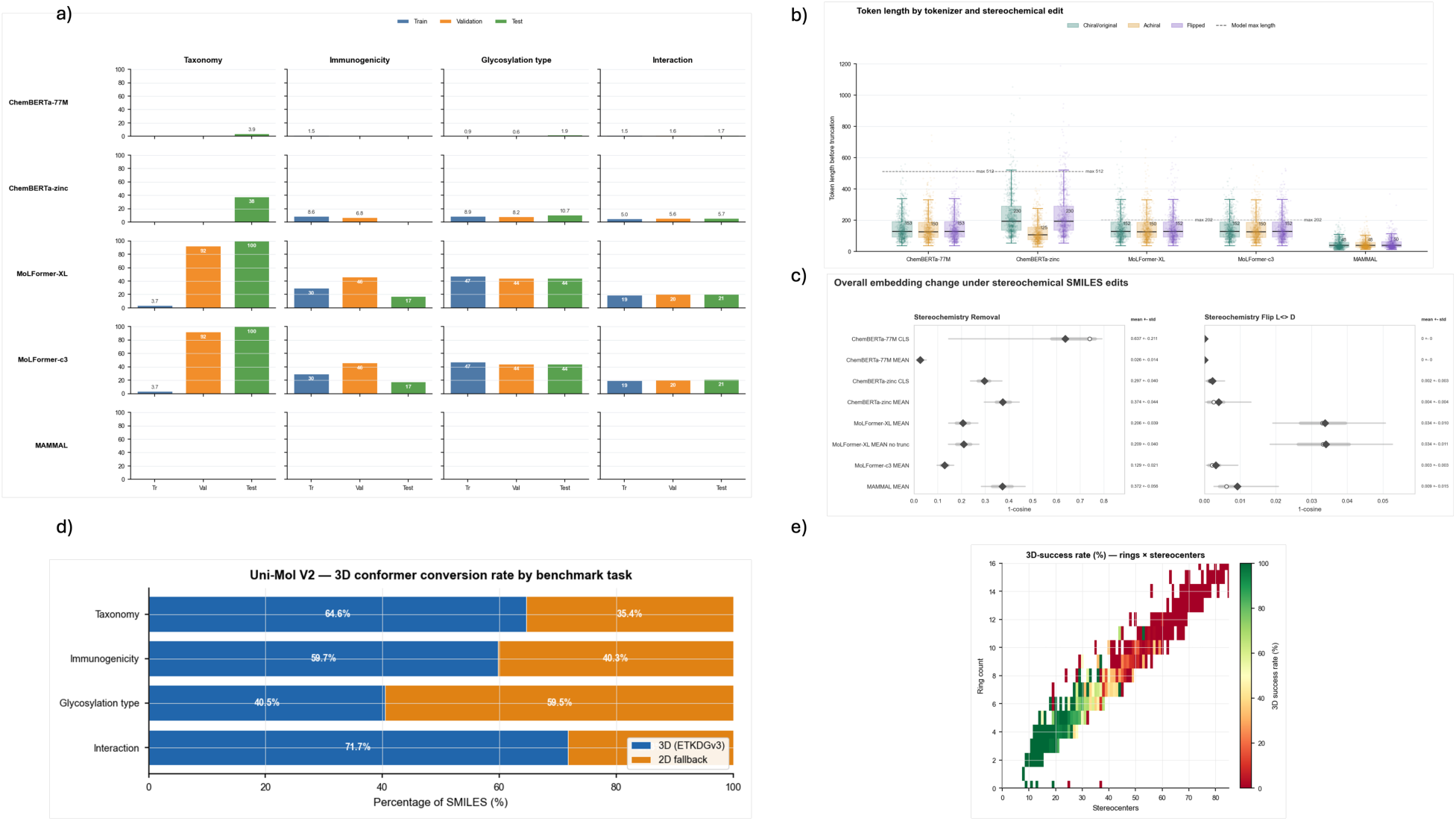
Molecular preprocessing and model architecture determine successful transfer to glycobiology. **a)** Percentage of glycans exceeding the effective input length of each SMILES-based foundation model after tokenization across benchmark tasks and dataset splits. MoLFormer showed extensive truncation, particularly for taxonomy datasets, whereas ChemBERTa retained most glycans. MAMMAL showed no truncation under the evaluated settings. **b)** Token-length distributions for chiral, achiral (stereochemistry removed) and stereochemically inverted SMILES before truncation. Dashed lines indicate each model’s pretrained context limit. Stereo removal substantially altered token lengths, whereas stereochemical inversion had minimal effects. **c)** Propagation of stereochemical perturbations into pretrained molecular representations. Left, cosine distances between embeddings extracted from matched chiral and achiral SMILES. Right, cosine distances following stereochemical inversion while preserving molecular connectivity. Stereo removal altered embeddings across all models, whereas stereochemical inversion revealed model-specific stereochemical sensitivity. **d)** Success of Uni-Mol V2 three-dimensional conformer generation across benchmark datasets. Bars show the proportion of glycans represented by ETKDG-generated conformers versus two-dimensional fallback structures. **e)** Relationship between glycan structural complexity and three-dimensional conformer generation. Heatmaps show conformer success as a function of ring count and stereocentre

Long-context tokenization preserves complex glycan structures. We first quantified sequence truncation after model-specific tokenization (Fig. 6a). Truncation varied markedly across models and benchmark tasks. ChemBERTa models, with effective input lengths of ∼512 tokens, retained most glycan structures, although ChemBERTa-zinc showed moderate truncation for the longest glycans. In contrast, MoLFormer, pretrained on canonical SMILES with molecules longer than 202 tokens excluded, exhibited extensive truncation, particularly for taxonomy benchmarks where validation and test glycans are substantially longer than the training set. In these datasets, approximately 92% of validation and 100% of test glycans exceeded MoLFormer’s input limit, indicating that many glycans were only partially represented. Functional datasets also showed substantial, although less severe, truncation. MAMMAL behaved differently. Its tokenizer generated much shorter glycan sequences and avoided truncation entirely, reflecting its multimodal tokenizer design. However, this did not translate into superior predictive performance, indicating that minimizing sequence length alone is insufficient if the resulting tokens do not preserve glycan-relevant chemistry.

Tokenization determines the molecular information presented to the encoder. Tokenized sequence lengths differed substantially between models (Fig. 6b), reflecting differences in tokenizer vocabularies and pretraining strategies. ChemBERTa-77M and MoLFormer produced similar median sequence lengths (∼129 tokens), ChemBERTa-zinc generated longer sequences (∼194 tokens), whereas MAMMAL produced much shorter sequences (∼38 tokens). Even within the ChemBERTa family, identical glycans were decomposed into different token sequences.

Stereochemical annotations further altered both tokenization and pretrained molecular representations (Fig. 6c; Supplementary Fig. 14). We previously identified stereochemistry as a key determinant of transferable glycan representations, with stereochemistry removal significantly reducing predictive performance across multiple benchmark tasks. To explain this effect mechanistically we examined whether the transition from chiral to achiral SMILES propagated through the representation-learning pipeline by comparing the cosine similarity between frozen embeddings extracted from matched chiral and achiral SMILES. Stereo removal changed token identities in all evaluated models and altered token sequence length in a model-dependent manner, with by far the largest token-length shifts observed for ChemBERTa-zinc.

These tokenizer-level changes propagated into the pretrained representation space, as stereochemistry removal also altered the resulting frozen molecular embeddings across all models. These results demonstrate that the effect of stereochemistry originates before downstream prediction, propagating from tokenizer-level changes into the pretrained molecular representation.

Stereochemistry removal and stereochemical inversion probe complementary aspects of molecular representation. Removing stereochemical annotations converts chiral SMILES into achiral SMILES, altering both stereochemical information and the resulting molecular string. We therefore performed a second, more controlled perturbation in which the stereochemical direction at every annotated stereocentre was inverted while preserving atom connectivity and molecular traversal. This transformation isolates the effect of stereochemical configuration without removing stereocentre configuration from the molecular representation. We quantified how this controlled inversion affected both model tokenization and the cosine similarity of the resulting frozen molecular embeddings (Supplementary Fig. 14). Whereas ChemBERTa-zinc, MoLFormer and MAMMAL generated distinct token sequences and corresponding embedding changes following stereochemical inversion, ChemBERTa-77M released tokenizer produced identical token sequences and therefore unchanged embeddings. Together, these analyses demonstrate that preserving stereochemistry requires two sequential properties: the tokenizer must first distinguish stereochemical configurations, and the pretrained encoder must subsequently propagate those differences into the molecular representation.

Finally, we assessed three-dimensional representation learning using Uni-Mol V2 (Fig. 6d,e). Successful conformer generation ranged from approximately 40-72% across benchmark datasets, with failed structures reverting to two-dimensional representations. Consequently, many nominally three-dimensional embeddings consisted of mixtures of genuine conformer-derived and fallback representations. Conformer generation was strongly dependent on glycan complexity. Glycans with fewer rings and stereocentres were converted successfully, whereas highly branched, stereochemically complex glycans, the structures of greatest biological interest, showed substantially lower success rates. Thus, current generic three-dimensional molecular pipelines remain constrained precisely where glycan structural complexity is greatest.

Collectively, these analyses explain why ChemBERTa-family models provide the strongest transferable glycan representations. MoLFormer was constrained by its effective 202-token pretrained context window, resulting in extensive truncation of large glycans, whereas current three-dimensional foundation models frequently failed to generate conformers for structurally complex glycans. MAMMAL avoided truncation through highly compressed tokenization but achieved lower predictive performance, indicating that compact molecular representations alone are insufficient for glycan modelling. In contrast, the longer context windows of ChemBERTa models preserved most glycan structures, allowing more complete molecular information to be presented during pretraining and subsequently retained in the extracted molecular embeddings.

Controlled stereochemical inversion further revealed that, unlike ChemBERTa-77M, ChemBERTa-zinc preserved stereochemical direction from tokenization into the resulting molecular representations. Together, these findings show that successful glycan transfer depends not only on foundation-model architecture, but also on how pretraining is constrained by tokenizer design, context length, and the propagation of stereochemical information into pretrained molecular representations.

### Chemical foundation models generalize across natural glycans and synthetic glycomimetics

Having established that ChemBERTa learns biologically meaningful glycan representations, we next asked whether these representations generalize beyond natural glycans to synthetic glycomimetics. We evaluated cross-domain transfer between CFG glycan-array interactions and glycomimetic binding measurements using protein concentration-aware protein-molecule representations (Fig. 7a). Protein sequences were encoded with ESM2, molecules with either ChemBERTa embeddings or ECFP6 Morgan fingerprints, and interaction strength was predicted using XGBoost.

**Figure 7.**
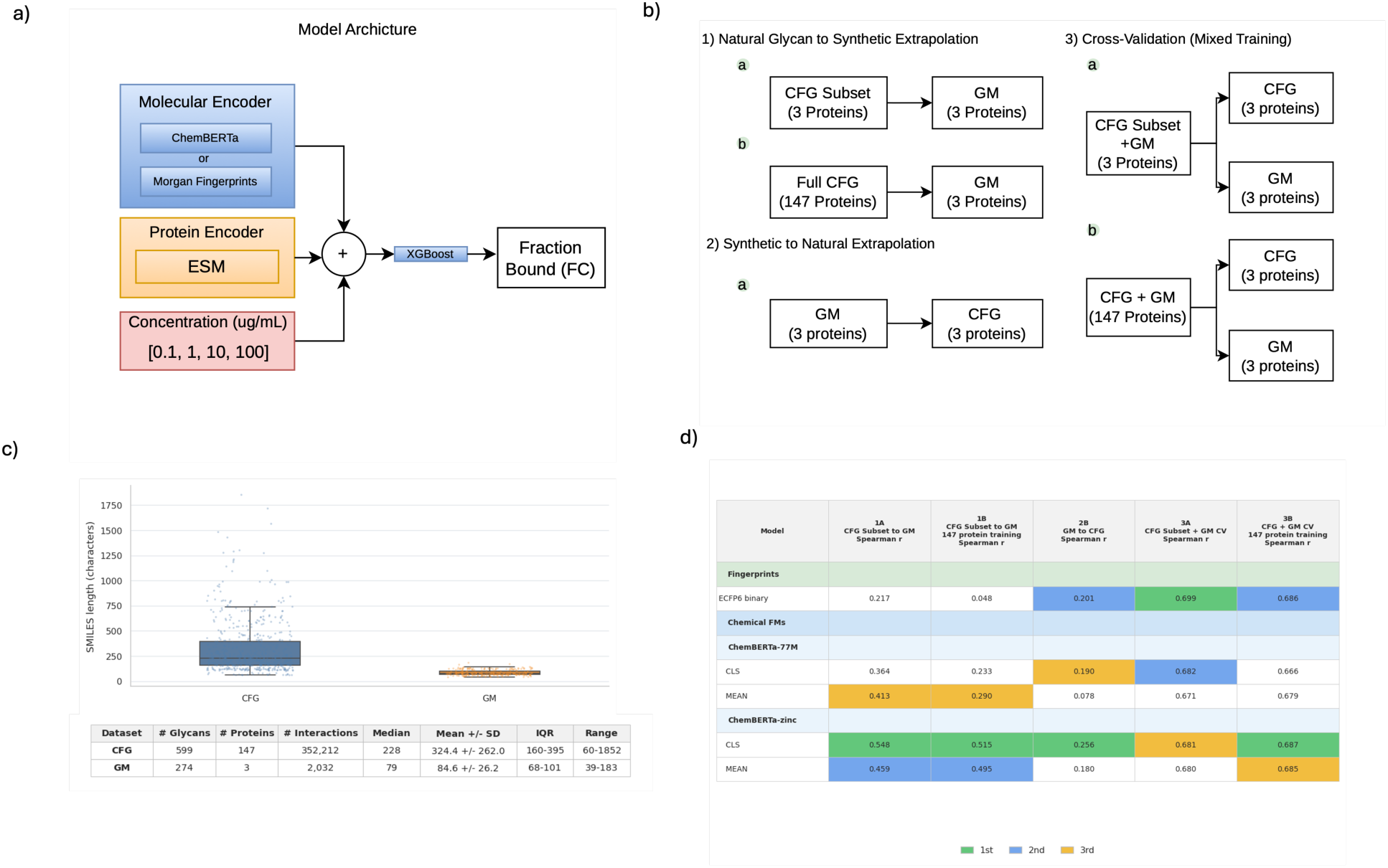
Chemical foundation models learn transferable molecular representations across natural glycans and synthetic glycomimetics. **a)** Protein-molecule interaction prediction framework. Protein sequences were encoded with ESM, molecules with either ChemBERTa embeddings or ECFP6 Morgan fingerprints, and protein concentration was included as an additional input. Combined features were used to predict fraction-bound interaction scores with XGBoost. **b)** Cross-domain evaluation strategy. Models were assessed using natural glycan-to-glycomimetic transfer, reciprocal glycomimetic-to-natural transfer, and mixed-domain training incorporating both molecular classes. **c)** Comparison of the CFG glycan-array and glycomimetic datasets. The CFG dataset comprised 599 glycans, 147 proteins and 352,212 interactions, whereas the glycomimetic dataset contained 274 molecules, three proteins and 2,032 interactions. Glycomimetics occupied a substantially smaller molecular-size regime than natural glycans. **d)** Prediction performance across transfer and mixed-domain settings. Bars show Spearman correlation between predicted and observed fraction-bound interaction scores. ChemBERTa-zinc achieved the strongest performance for direct cross-domain transfer, whereas ChemBERTa and ECFP6 performed similarly when natural glycans and glycomimetics were jointly represented during training.

The two datasets differed substantially in scale and molecular complexity (Fig. 7b,c). The CFG dataset comprised 599 natural glycans measured against 147 proteins (352,212 interactions), whereas the glycomimetic dataset contained 274 synthetic molecules measured against three shared galectins (2,032 interactions). Glycomimetics were also considerably smaller than natural glycans, occupying a chemical regime closer to conventional small molecules and therefore providing a stringent test of cross-domain transfer. We evaluated direct natural-to-synthetic and synthetic-to-natural extrapolation together with mixed-domain learning (Fig. 7b).

ChemBERTa models consistently outperformed Morgan fingerprints in the direct transfer settings (Fig. 7d). Training on natural glycans and testing on glycomimetics, ChemBERTa-zinc CLS achieved the strongest performance (Spearman *r* = 0.548 versus 0.217 for ECFP6), and retained a substantial advantage when trained on the full 147-protein CFG panel (*r* = 0.515 versus 0.048). Reciprocal glycomimetic-to-CFG transfer also favoured ChemBERTa (*r* = 0.256 versus 0.201). By contrast, when natural glycans and glycomimetics were jointly represented during training, ChemBERTa and ECFP6 performed similarly, with fingerprints remaining highly competitive in the best-represented mixed-domain setting.

These findings demonstrate that ChemBERTa captures transferable molecular features that generalize across chemically distinct regions of carbohydrate space. Rather than simply memorizing glycan-specific patterns, the model learns atom-level determinants of molecular recognition that remain informative when transferred between natural glycans and structurally simplified synthetic glycomimetics, supporting the use of chemical foundation models for glycomimetic discovery and design.

### Chemical foundation models recover experimentally validated glycan-lectin recognition

Having shown that ChemBERTa representations transfer from natural glycans to synthetic glycomimetics, and that stereochemical information is differentially preserved from tokenization into pretrained molecular embeddings, we next asked whether these representations recover independent experimental measurements of glycan-lectin recognition. We evaluated the chiral molecular representations from the 147-protein mixed-domain interaction models on the independent Liquid Glycan Array (LiGA), without retraining on LiGA labels. LiGA provides a particularly stringent validation because it contains a targeted panel of closely related, stereochemically defined glycans, including matched D/L stereoisomers with identical molecular composition and connectivity but different three-dimensional configurations, represented by small changes in their stereochemical SMILES annotations (*see Methods*). The panel comprised 66 interactions between 11 lectins and six glycans, of which 15 were experimentally validated binders.

Mean-pooled ChemBERTa representations provided the strongest prioritization of experimentally validated binders. Across the four individual concentration inputs, ChemBERTa-77M MEAN achieved the highest mean AP across protein concentrations (AP = 0.391), followed by ChemBERTa-zinc MEAN (AP = 0.355), with both representations showing stronger binder prioritization at higher concentration inputs (Fig. 8a; Supplementary Fig. 15 and 16). When predictions were first averaged across concentrations, ChemBERTa-zinc MEAN achieved the highest AP (0.403), followed by ChemBERTa-77M MEAN (0.387); both outperformed the random expectation and the ECFP6 fingerprint baseline (AP = 0.264) (Fig. 8b,c). ChemBERTa-77M MEAN produced the strongest overall ordering of positive relative to non-positive interactions (AUROC = 0.678; nominal one-sided Mann–Whitney U P = 0.019), whereas ChemBERTa-zinc MEAN achieved an AUROC of 0.631 (nominal P = 0.063). These results demonstrate that molecular representations learned during general chemical pretraining transfer to an independent experimental assay and successfully prioritize experimentally validated glycan-lectin interactions.

**Figure 8.**
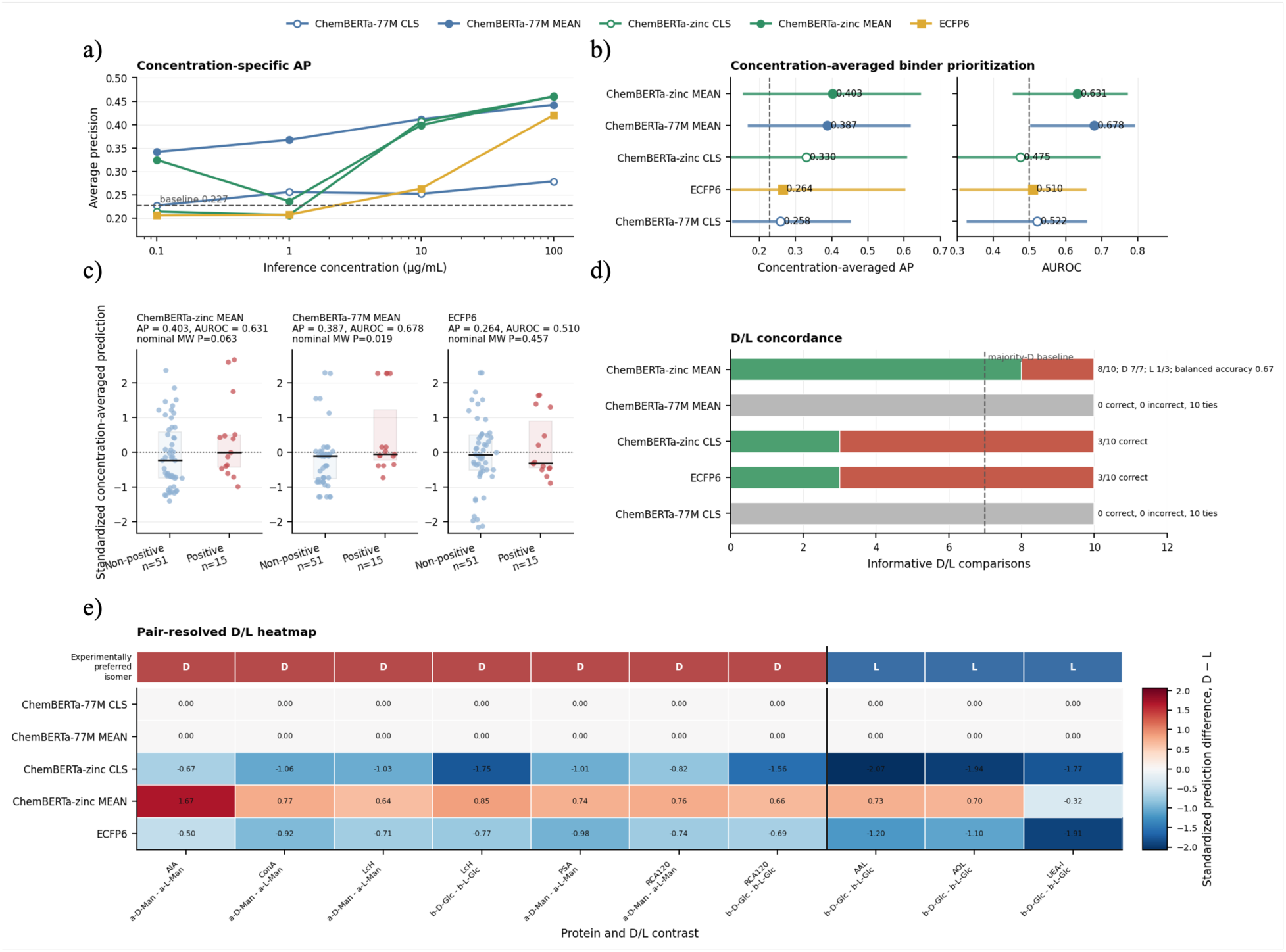
Independent LiGA validation separates binder prioritization from stereochemical discrimination. **a)** Average precision (AP) for prioritizing experimentally validated LiGA glycan-lectin interactions at protein concentrations of 0.1, 1, 10 and 100 µg/mL. AP was calculated using all 66 lectin-glycan pairs. Dashed line indicates the prevalence baseline (15/66 = 0.227). **b)** Concentration-averaged binder prioritization. Predictions were averaged across concentrations for each lectin-glycan pair. Left, AP; right, AUROC. Dashed lines indicate random expectation. Error bars show lectin-cluster bootstrap 95% confidence intervals. **c)** Concentration-averaged standardized prediction scores for experimentally positive (n = 15) and non-positive (n = 51) interactions. Points represent individual pairs; boxes show interquartile ranges and medians. *P* values were calculated using one-sided Mann-Whitney tests. **d)** Recovery of experimentally observed D/L stereochemical preference across 10 matched stereoisomer pairs. Predictions were averaged across concentrations before comparing experimentally positive and non-positive stereoisomers. Bars show correct, incorrect and tied predictions. Dashed line indicates the majority-D baseline (7/10). **e)** Pairwise D/L stereochemical preference. Heatmap shows standardized D/L prediction differences for the same 10 stereoisomer pairs. Positive values favour the D isomer and negative values the L isomer. ChemBERTa-zinc MEAN showed the strongest stereochemical concordance, whereas ChemBERTa-77M was invariant to D/L configuration.

Binder prioritization and stereochemical discrimination emerged as distinct model capabilities. We therefore examined ten informative D/L contrasts in which one stereoisomer was experimentally positive, and its matched counterpart was non-positive (Fig. 8d,e). ChemBERTa-77M CLS and MEAN assigned identical scores to every D/L pair, producing ten ties (as expected from mechanistic analysis). In contrast, ChemBERTa-zinc MEAN representation assigned the higher score to the experimentally preferred stereoisomer in 8 of 10 comparisons, whereas ChemBERTa-zinc CLS and ECFP6 each recovered 3 of 10.

The D/L panel contained seven D-positive and three L-positive contrasts, and ChemBERTa-zinc MEAN exhibited an overall preference for D-configured glycans. It correctly recovered all seven D-positive comparisons and one of the three L-positive comparisons, corresponding to a balanced accuracy of 0.67. Its 8 out of 10 concordance therefore represents the strongest stereochemical recovery among the evaluated model representations, but remains influenced by a global D-isomer preference. These findings motivate the development of larger, balanced LiGA panels containing comparable numbers of D- and L-selective glycan-protein interactions to enable more rigorous evaluation of lectin-specific stereochemical recognition.

Together, the independent LiGA analysis separates two capabilities of molecular foundation models. ChemBERTa-77M MEAN retained broad interaction-relevant chemical information and effectively prioritized experimentally validated binders despite being invariant to stereochemical direction. ChemBERTa-zinc, by contrast, preserved configurational information from tokenization through pretrained molecular embeddings into differential interaction predictions, and its mean-pooled representation provided the strongest combined recovery of binder prioritization and D/L preference.

Together with the natural-to-synthetic transfer results, these findings provide independent experimental evidence that chemical foundation models capture transferable molecular determinants of carbohydrate recognition. By tracing stereochemical information from molecular tokenization and general chemical pretraining through latent representations to experimentally measured lectin binding, this study establishes a mechanistic framework for prioritizing glycan-protein interactions and designing stereochemically informed glycomimetic ligands.

## Discussion

Chemical foundation models have transformed molecular machine learning, yet glycans have generally been regarded as requiring specialised molecular representations because of their branched architectures, stereochemical complexity and non-linear biosynthesis. Here, we show that representations learned from general molecular chemistry transfer effectively to glycobiology, and the extent of transfer depends strongly on both the biological task and the underlying model architecture. Glycan-native representations remained optimal for taxonomy, where explicit encoding of glycan syntax and biosynthetic hierarchy is advantageous, whereas transferable atom-level representations proved most effective for molecular-recognition tasks, including immunogenicity and protein-glycan interaction prediction.

A central finding of this study is that biological supervision reorganizes, rather than replaces, transferable chemical representations. Pretrained ChemBERTa embeddings already contained chemically coherent organisation reflecting shared monosaccharide composition, linkage chemistry and glycan architecture despite no explicit knowledge of glycosylation pathways or glycan grammar. Fine-tuning progressively reorganized these latent chemical neighbourhoods into biologically meaningful programmes associated with biosynthesis, immunogenicity, molecular recognition and evolutionary origin. Depending on the supervisory signal, the same underlying chemical representation emphasized canonical N- and O-glycan biosynthetic pathways, separated mammalian from microbial and plant-associated carbohydrate chemistries, or grouped glycans according to shared recognition by lectins, microbial adhesins and viral haemagglutinins. These observations indicate that many biologically relevant dimensions of glycobiology are already embedded within general chemical representations and can be selectively exposed through task-specific supervision.

Mechanistic analyses identified four properties governing successful transfer to glycobiology: stereochemical preservation, tokenizer strategy, context length and complete representation of the glycan structure. Rather than model size alone, these architectural features largely explained performance differences between foundation models. ChemBERTa-zinc preserved stereochemical information while retaining sufficient molecular context for large branched glycans. In contrast, MoLFormer models were more strongly affected by sequence truncation, ChemBERTa-77M exhibited reduced sensitivity to stereochemical directionality, MAMMAL compressed molecular information more aggressively, and current three-dimensional models were limited by incomplete conformer generation and atom-count constraints. These findings suggest that successful transfer depends primarily on preserving the molecular information most relevant to carbohydrate chemistry rather than increasing model scale.

The natural-to-synthetic transfer experiments further demonstrate that these representations capture molecular-recognition principles extending beyond naturally evolved glycans. Models trained on natural glycan arrays generalized to synthetic glycomimetics, while reciprocal transfer retained predictive signal despite substantial differences in scaffold composition and chemical substitution. This capability establishes a computational bridge between glycobiology and glycomimetic chemistry, suggesting that representations learned from biological carbohydrates can be extended to chemically engineered ligands and providing a foundation for foundation-model-guided glycomimetic discovery.

Independent LiGA validation provided experimental evidence that these transferable representations recover biologically meaningful stereochemical recognition. ChemBERTa consistently outperformed fixed molecular fingerprints in prioritizing experimentally validated glycan-lectin interactions and more accurately recovered D/L-selective binding preferences, indicating that the learned representations capture atom-level determinants of carbohydrate recognition rather than broad glycan-family associations. Together with the cross-domain transfer experiments, these results demonstrate that transferable chemical representations retain experimentally relevant molecular information across independent glycan recognition platforms.

Several limitations should be acknowledged. Immunogenicity labels represent sequence-level annotations and do not capture additional determinants of immune recognition, including molecular presentation, epitope density, carrier context or host-specific responses. GlyLES conversion excluded a subset of glycans containing unsupported residues or modifications, while the LiGA validation panel comprised a relatively small number of lectins and stereochemically informative interactions. Furthermore, LiGA provides semi-quantitative interaction annotations rather than quantitative affinity measurements, preventing direct assessment of concentration-dependent predictions. Larger experimentally validated datasets encompassing broader glycan diversity and quantitative binding measurements will be important for establishing the generality of these findings.

More broadly, our results suggest that glycans need not be viewed as a computational domain entirely separate from general molecular chemistry. Instead, transferable chemical representations and glycan-native representations provide complementary descriptions of glycan biology. Glycan-native encodings remain advantageous for structurally defined tasks closely linked to glycan biosynthesis and taxonomy, whereas atom-level chemical representations are particularly effective for molecular-recognition problems and for transferring knowledge between natural and synthetic carbohydrate chemistry. This complementary view is likely to be more productive than treating the two representation paradigms as competing alternatives.

Overall, this work demonstrates that general chemical foundation models already encode transferable molecular information relevant to glycobiology and that biological supervision progressively specializes these representations toward distinct biological functions. By combining comparative benchmarking, mechanistic analyses of representation learning, cross-domain transfer and independent experimental validation, we show that glycobiological organisation can emerge from representations learned from general molecular chemistry. These findings provide a foundation for integrating advances in molecular foundation models with glycobiology, enabling improved prediction of glycan function, protein-glycan recognition and the rational design of next-generation glycomimetics.

## Methods

### GlycanMLBenchmark datasets and split strategy

We evaluated chemical foundation models (ChemFMs) using GlycanMLBenchmark^34^, a standardized multi-task benchmark dataset including 11 downstream tasks across 4 different areas of glycobiology: taxonomy prediction, immunogenicity prediction, glycosylation type prediction and protein-glycan interaction prediction. Unless otherwise stated, the original benchmark training, validation and test splits were retained.

**Taxonomy prediction**, derived from SugarBase^29^, consists of eight hierarchical classification tasks corresponding to domain, kingdom, phylum, class, order, family, genus and species. The original benchmark employs a motif-based partitioning strategy in which glycans are first clustered by K-means (K = 10), followed by assignment of eight clusters to the training set and one cluster each to the validation and test sets. During dataset curation, we identified taxonomic labels present in the validation or test partitions that were absent from the corresponding training partition, rendering these classes unlearnable under supervised learning. To ensure valid model training and evaluation, taxonomy datasets were filtered so that every label present in the validation and test sets was represented in the training data. In addition, because some glycans were associated with multiple taxonomic annotations whereas the benchmark defines taxonomy as a single-label classification task, a single annotation was retained for each glycan by selecting the first recorded label. Model performance was evaluated using the macro-averaged F1 score (Macro-F1).

**Immunogenicity Prediction**, also derived from SugarBase^29^, was formulated as a binary classification task to predict whether a glycan is immunogenic or non-immunogenic. Dataset partitioning followed the original motif-based 8:1:1 split generated from the subset of glycans with immunogenicity annotations. Performance was evaluated using area under the precision-recall curve (AUPRC).

**Glycosylation type prediction,** derived from GlyConnect ^46^, was formulated as a classification task predicting whether a glycan is N-glycosylated, O-glycosylated, or free. The original splitting strategy was also the motif-based partitioning strategy (8:1:1). Performance was evaluated on Macro-F1.

**Protein-glycan interaction prediction** was derived from LectinOracle^27^, which was curated from the Consortium of Functional Glycomics (CFG) ^47^. Unlike the glycan-only tasks, this benchmark was formulated as a regression problem to predict protein-glycan binding affinity, represented as the Z-score of transformed relative fluorescence units. To minimise information leakage arising from homologous proteins, dataset partitioning was performed according to protein sequence similarity rather than glycan structure. Protein sequences were clustered using MMseqs2^48^ with a minimum within-cluster sequence identity of 50%, after which clusters were assigned to the training, validation and test sets in an 8:1:1 ratio. Individual protein-glycan interaction pairs were then allocated according to the corresponding protein partition. Model performance was evaluated using Spearman’s rank correlation coefficient (Spearman’s ρ).

### Glycan processing and molecular representation generation

Benchmark glycans were provided as IUPAC-condensed sequences, which were converted into three complementary molecular representations for downstream modelling. Sequence representations consisted of IUPAC-derived tokens encoding individual monosaccharides and glycosidic linkages. Graph representations modelled glycans as molecular graphs in which nodes represented monosaccharides and edges represented glycosidic linkages. Engineered graph descriptors included branching, connectivity, centrality, core-corona organisation and node-type features. SMILES representations were generated from IUPAC-condensed sequences using GlyLES. Glycans containing unsupported monosaccharides or structural modifications that could not be converted by GlyLES were excluded from all subsequent analyses (Supplementary Table 1). To ensure direct comparability across representation families, all benchmark comparisons were performed on the same GlyLES-convertible subset of glycans.

### Molecular representation extraction and downstream modelling

We compared fixed molecular representations and task-adaptive representation-learning approaches to evaluate the transferability of chemical foundation models across glycobiological prediction tasks. Fixed representations comprised Morgan fingerprints, engineered glycan graph descriptors and frozen embeddings extracted from pretrained chemical foundation models. Chiral SMILES and chirality-aware Morgan fingerprints were used throughout the primary analyses. Morgan fingerprints were generated using RDKit as extended-connectivity fingerprints with radii of two (ECFP4) and three (ECFP6), and both binary and count-based fingerprint variants were evaluated.

For SMILES-based chemical foundation models, glycan SMILES were tokenized using the native tokenizer associated with each pretrained model and encoded according to the recommended embedding strategy for that architecture. We evaluated two ChemBERTa-family models (ChemBERTa-ZINC and ChemBERTa-77M), two MoLFormer-family models (MoLFormer-XL-both-10pct and MoLFormer-1.1B^49^), MAMMAL, and Uni-Mol V2. Where supported, both CLS-token representations and mean-pooled token embeddings were extracted. Tokenized glycan SMILES were processed using the effective input-length constraints of the corresponding pretrained model implementation.

For MoLFormer-family models, we additionally evaluated a no-truncation configuration, in which truncation was disabled during embedding extraction. Uni-Mol V2 three-dimensional molecular inputs were generated from glycan SMILES using the default RDKit ETKDG conformer generation procedure used by the Uni-Mol V2 pipeline. When ETKDG conformer generation failed, the corresponding molecule was retained using the two-dimensional fallback representation generated from the molecular graph. Detailed model architectures, tokenizer specifications and pretraining characteristics are provided in Table 4.

### Static representation benchmarking

For the static representation benchmark, pretrained molecular representations remained fixed and only the downstream prediction model was trained. Fixed glycan representations were used as input to lightweight downstream predictors, including XGBoost and shallow multilayer perceptron heads. Classification heads were used for taxonomy, immunogenicity and glycosylation-type prediction, whereas regression heads were used for protein-glycan interaction prediction. For protein-glycan interaction prediction, protein sequences were encoded using frozen ESM2 embeddings by mean pooling over token-level protein representations. Protein and glycan embeddings were concatenated before downstream regression. The protein encoder remained fixed throughout all experiments, allowing model performance to reflect differences in glycan molecular representation rather than changes in protein encoding.

Neural downstream heads were optimised using task-specific objective functions: cross-entropy for taxonomy and glycosylation-type prediction, binary cross-entropy with logits for immunogenicity prediction, and mean squared error for protein-glycan interaction prediction. Early stopping based on validation performance was applied to all neural models.

For the task-adaptive setting, we compared two trainable representation-learning strategies. First, we evaluated glycan-specialist models trained directly on glycan-native representations. Sequence-based architectures included convolutional neural networks (CNNs), ResNet, long short-term memory (LSTM) networks and bidirectional LSTMs trained on IUPAC-derived sequence tokens. Graph-based architectures included graph attention networks (GAT), graph isomorphism networks (GIN), message-passing neural networks (MPNN), relational graph convolutional networks (RGCN) and CompGCN operating on glycan graph representations. These models were trained from scratch for each downstream task without chemical pretraining using the architectures and hyperparameter settings defined in the GlycanMLBenchmark repository. Results were averaged across three independent random seeds.

Second, we fine-tuned pretrained chemical foundation models using glycan SMILES as input. Task-specific prediction heads were attached to each molecular encoder and the complete glycan branch was optimized end-to-end. For protein-glycan interaction prediction, the ESM2 protein encoder remained frozen, while the glycan encoder and downstream prediction head were updated. Models were trained using the same task-specific objective functions as the static benchmark and selected according to validation performance using early stopping. To directly compare pretrained and fine-tuned representations under identical downstream conditions, embeddings were extracted from the best-performing validation checkpoint following fine-tuning and evaluated using the same XGBoost and MLP prediction pipeline employed for frozen representations. This ensured that differences in downstream performance reflected changes in the learned molecular representation rather than differences in the prediction model. Complete hyperparameter settings for all experiments are provided in Supplementary Table 6.

### Embedding-space clustering and motif enrichment analysis

For each benchmark task, embedding-space analyses were performed using the model and pooling strategy selected by post-fine-tuning validation performance. Pretrained and fine-tuned embeddings from the selected configuration were then analysed separately to assess how task-specific adaptation altered the organization of the learned glycan representation space. Embeddings were visualized using t-SNE with cosine distance. Pre- and post-fine-tuning embeddings were projected independently, rather than fitted jointly, so that each plot reflected the geometry of the embedding space at that stage. The t-SNE projections were used only for visualization; cluster assignments were not derived from the two-dimensional coordinates.

Biological communities (clusters) were identified directly from the original high-dimensional embeddings. For each representation space, a cosine similarity neighbourhood graph was constructed and partitioned using the Leiden community detection algorithm. Because the optimal community structure differed between downstream tasks, the number of clusters was not specified a priori but determined independently for each embedding space through a systematic resolution-selection procedure.

Leiden resolution parameters were evaluated across a predefined resolution sweep. At each candidate resolution, clustering quality was quantified using the average silhouette score calculated with cosine distance on the original embedding vectors. Cluster stability was assessed by calculating the adjusted Rand index (ARI) between neighbouring resolutions and across multiple Leiden random initializations. Resolutions generating highly fragmented communities or unstable clustering solutions were excluded. The final resolution was selected as the lowest value whose silhouette score was within 0.02 of the maximum observed value while satisfying the stability and minimum cluster-size criteria. The resulting Leiden communities defined the embedding-space partitions used for all subsequent analyses.

To interpret the biological organisation of the learned representations, each Leiden community was analysed using cluster-versus-rest motif enrichment analysis implemented in Glycowork. Glycans assigned to an individual community were compared with all remaining glycans, and significantly enriched glycan motifs were identified following multiple-testing correction. Enriched motifs were subsequently integrated with the benchmark annotations to determine whether individual embedding communities were preferentially associated with specific biological labels before or after fine-tuning. This analysis enabled changes in embedding-space organisation to be interpreted in terms of biologically meaningful glycan motifs and functional recognition programmes.

### Stereochemistry Ablation and Analysis

To assess the contribution of stereochemical information to glycan representation learning, we repeated the static benchmark using matched SMILES inputs in which stereochemical annotations were removed. Stereochemical SMILES markers were stripped from the encoder-facing SMILES strings while keeping glycan identities, labels, and train-test splits unchanged. Embeddings were then re-extracted and evaluated using the same downstream benchmark models and metrics as the original representations. For fingerprint baselines, matched chiral and achiral Morgan fingerprints were compared using the same radius and fingerprint size. The effect of stereochemistry was quantified as the paired performance difference between the original or chiral representation and its no-chirality or achiral counterpart. Because each representation had a matched stereochemistry-preserving and stereochemistry-removed version, statistical significance was assessed using paired tests. Performance differences were first standardized within each grouped benchmark task to make effects comparable across metrics. We then applied exact two-sided Wilcoxon signed-rank tests to the paired standardized differences, testing whether the median effect of stereochemistry differed from zero. Multiple testing was controlled using the Benjamini-Hochberg false discovery rate procedure within each model subset, and rank-biserial correlation was reported as the paired non-parametric effect size.

### Tokenization, truncation and conformer-generation analysis

We examined model-specific SMILES tokenization, stereochemical handling, sequence truncation and three-dimensional conformer generation. Glycan SMILES were processed using each model’s native tokenizer. ChemBERTa-zinc uses a ByteLevel tokenizer and ChemBERTa-77M its checkpoint-specific molecular RoBERTa tokenizer; both were evaluated with a maximum input length of 512 tokens. MoLFormer uses a regex-based SMILES tokenizer. Because molecules longer than 202 tokens were excluded from MoLFormer pretraining, 202 tokens were treated as its effective pretrained context length. A no-truncation configuration was additionally evaluated as an out-of-distribution diagnostic. MAMMAL inputs were encoded using its modular molecular prompt tokenizer, with no truncation observed under the tested settings. Two matched stereochemical perturbations were generated for each glycan. For stereochemistry removal, RDKit was used to generate a canonical non-isomeric SMILES. For controlled stereochemical inversion, all @ and @@ direction tags were swapped while preserving atom traversal, connectivity and all non-stereochemical characters. This operation tested sensitivity to stereochemical direction and was not assumed to represent a formal D-to-L conversion for every complex glycan. Original, stereo-removed and inverted SMILES were tokenized before truncation.

We recorded token IDs and sequence lengths, calculated the proportion of molecules whose token sequences changed, and measured the corresponding token-length differences. Frozen embeddings were then extracted using the same pooling strategies as in the static benchmark. Representation changes were quantified as cosine distance between the original and perturbed embeddings. Because embedding geometries differ across models, cosine-distance magnitudes were interpreted within each representation rather than compared directly between models. For truncation analysis, pre-truncation token lengths were compared with each model’s effective context limit. Uni-Mol V2 was analysed separately because it uses molecular graphs and atomic coordinates rather than SMILES tokens. We recorded whether preprocessing generated a valid three-dimensional conformer or required a two-dimensional fallback, and whether molecules exceeded the 128-atom input limit. Conformer-generation success was further related to RDKit-derived molecular descriptors, including heavy-atom, ring, stereocentre and rotatable-bond counts, molecular weight and SMILES length.

### Natural glycan-glycomimetic transfer learning

To evaluate whether chemical foundation models learned transferable molecular representations beyond naturally occurring glycans, we performed cross-domain prediction experiments using the benchmark introduced by *Carpenter et al* ^33^. This benchmark integrates natural glycan array measurements from the Consortium for Functional Glycomics (CFG) with experimentally measured binding data for synthetic glycomimetics, enabling direct evaluation of transfer learning between natural and synthetic carbohydrate chemistries.

The natural glycan component comprised 599 glycans profiled against 147 glycan-binding proteins at four protein concentrations, yielding 352,212 protein-glycan interaction measurements. The synthetic component comprised 274 glycomimetics evaluated against the three galectins shared with the CFG dataset (Galectin-1, Galectin-3 and Galectin-7), generating 2,032 protein-glycomimetic interaction measurements.

### Response harmonisation

The original CFG measurements were reported as relative fluorescence units (RFU). Following the *Carpenter et al*. processing framework, RFU values were background-corrected, averaged across replicate array measurements and normalized to the assay response range to obtain fraction-bound values constrained between 0 and 1. Glycomimetic binding measurements were extracted from the accompanying publication and supplementary tables. Where glycomimetic binding was reported through affinity measurements, including association constants, these values were converted to the same fraction-bound response scale using the concentration-dependent binding formulation used in the source study. This harmonisation produced a common quantitative response variable spanning both natural glycans and synthetic glycomimetics. Protein concentrations were similarly standardised across datasets. All interaction measurements were mapped to a common four-point concentration grid comprising 0.1, 1, 10 and 100 µg/mL allowing the two experimental domains to be modelled using a unified regression framework.

### Molecular representation and prediction model

For each interaction example, the input vector consisted of a molecular representation, a protein representation and the protein concentration feature. Molecules were represented using either ChemBERTa embeddings extracted from glycan or glycomimetic SMILES, or ECFP6 Morgan fingerprints. Protein sequences were represented using frozen ESM embeddings obtained by mean pooling over residue-level protein representations. Molecular embeddings, protein embeddings and the protein concentration feature were concatenated and used as input to an XGBoost regression model trained to predict fraction-bound binding. Predictive performance was evaluated using Spearman’s rank correlation coefficient (Spearman’s ρ).

### Transfer-learning evaluation

We evaluated five transfer settings matching the *Carpenter et al*. benchmark design. In setting 1a, models were trained on CFG interactions for the three proteins shared with the glycomimetic dataset. Ten-fold cross-validation was performed on the CFG subset containing 7,188 observations, and a final model was then trained on all CFG rows from this subset and evaluated on the held-out glycomimetic domain containing 2,032 observations. This setting directly tested CFG-to-glycomimetic extrapolation for the same three galectins.

In setting 1b, models were trained on the full CFG protein panel. Ten-fold cross-validation was performed on 352,212 CFG observations from 147 proteins, after which a final model was trained on all CFG rows and evaluated on the same held-out glycomimetic set of 2,032 observations for Galectin-1, Galectin-3 and Galectin-7. This tested whether broader CFG protein diversity improved transfer to glycomimetics.

In setting 2a, the transfer direction was reversed. Models were trained using the 2,032 glycomimetic observations across the three shared galectins, with ten-fold cross-validation performed within the glycomimetic domain. A final model was then trained on all glycomimetic rows and evaluated on the held-out CFG subset for the same three galectins, comprising 7,188 observations. This setting tested glycomimetic-to-natural-glycan extrapolation.

Settings 3a and 3b assessed mixed-domain learning. In setting 3a, CFG and glycomimetic observations were combined for the three shared galectins (Galectin-1, Galectin-3 and Galectin-7) giving 9,220 observations in total. In setting 3b, the glycomimetic observations were combined with the full 147-protein CFG dataset, giving 354,244 observations in total. Both settings were evaluated using the original ten-fold molecular-level cross-validation scheme, with all observations associated with the same molecule, including measurements across concentrations, retained within the same fold. Performance was calculated from pooled out-of-fold predictions across all ten folds. For the extrapolation settings involving an external target domain, transfer performance was evaluated on the held-out target domain after retraining the final model on all available source-domain observations.

### Independent experimental validation using liquid glycan arrays (LiGA)

To independently evaluate whether the learned molecular representations captured experimentally validated glycan-lectin recognition, we tested the interaction models on the Liquid Glycan Array (LiGA) validation dataset reported by Carpenter et al.²¹. This dataset was not used during model training and therefore provided an independent assay-level experimental assessment of model generalisation. The LiGA validation panel comprised 11 lectins (UEA-1, AAL, LcH, RCA120, AIA, AOL, ConA, Galectin-1, Galectin-3, Galectin-7 and PSA) tested against six stereochemically defined monosaccharides (α-D-Man, α-L-Man, β-D-Glc, β-L-Glc, α-L-Glc and α-L-Rha). The resulting dataset contained 66 lectin–glycan pairs, of which 15 were annotated as experimentally positive binding interactions after confirmation in clonal assays, while the remaining 51 pairs were treated as non-positive interactions.

### Prediction pipeline

For each LiGA glycan, the corresponding SMILES representation was used to compute the same molecular features used in the natural glycan-glycomimetic transfer experiments. We focused on the two representation families evaluated in the LiGA validation analysis: ChemBERTa (ChemBERTa-ZINC and ChemBERTa-77M) molecular embeddings and ECFP6 Morgan fingerprints. Lectin sequences were encoded using the same frozen ESM protein-embedding procedure used in the transfer experiments, obtained by mean pooling residue-level protein representations. Molecular features, ESM protein embeddings and concentration values were concatenated and passed to the trained XGBoost interaction models to generate predicted fraction-bound scores. LiGA predictions were generated using the mixed-domain interaction models previously trained on combined CFG natural glycan and glycomimetic data, 3b setting. Each LiGA lectin-glycan pair was evaluated at the same four concentration inputs used in the transfer experiments: 0.1, 1, 10 and 100 µg/mL. For each model-concentration combination, predicted scores were evaluated. Model performance was evaluated using average precision as the primary ranking metric because the validation set was class imbalanced. To assess whether experimentally positive interactions received systematically higher predicted scores than non-positive interactions, predicted scores were standardized within each model–concentration combination and compared using one-sided Mann–Whitney U tests. Finally, we evaluated whether each model recovered stereochemistry-dependent binding preferences. For each informative D/L monosaccharide contrast, model predictions were compared between the experimentally binding stereoisomer and its matched non-binding counterpart. A contrast was counted as correctly recovered when the experimentally positive stereoisomer received the higher predicted score. This analysis was used to quantify whether the interaction model preserved stereochemical specificity in glycan-lectin recognition.

## Data Availability

All datasets generated or analysed during this study, including the processed GlycanMLBenchmark datasets, GlyLES-converted SMILES representations, benchmark splits and all associated datasets and metadata required to reproduce the reported analyses, will be made publicly available upon publication.

## Code Availability

All code used for dataset preprocessing, benchmark evaluation, embedding extraction, model training and fine-tuning, statistical analyses, representation analyses and figure generation will be made publicly available upon publication

## Author Contributions

M.S.R., J.B.S. and G.P.D. conceived and designed the study. M.S.R., J.B.S. and G.P.D. developed the methodology, designed the analyses, interpreted the results and contributed to the overall scientific direction of the work. M.S.R.performed the computational experiments, data processing, model development, statistical analyses and visualization, and prepared the original manuscript. M.S.R., J.B.S. and G.P.D. contributed to writing, reviewing and editing the manuscript. V.L.G., R.F.D. and N.M. contributed to scientific discussion, interpretation of the results, and critical review and editing of the manuscript. All authors reviewed and approved the final manuscript.

## Use of generative artificial intelligence

Generative artificial intelligence tools were used during this study to assist with language editing and refinement of the manuscript, as well as with code development, debugging and documentation. All AI-assisted text and code were reviewed, edited and verified by the authors, who take full responsibility for the accuracy, integrity and reproducibility of the work. Scientific interpretation, analysis of results and conclusions were determined by the authors.

## Supplementary Material

### Supplementary Figures

**Supplementary Figure 1.**
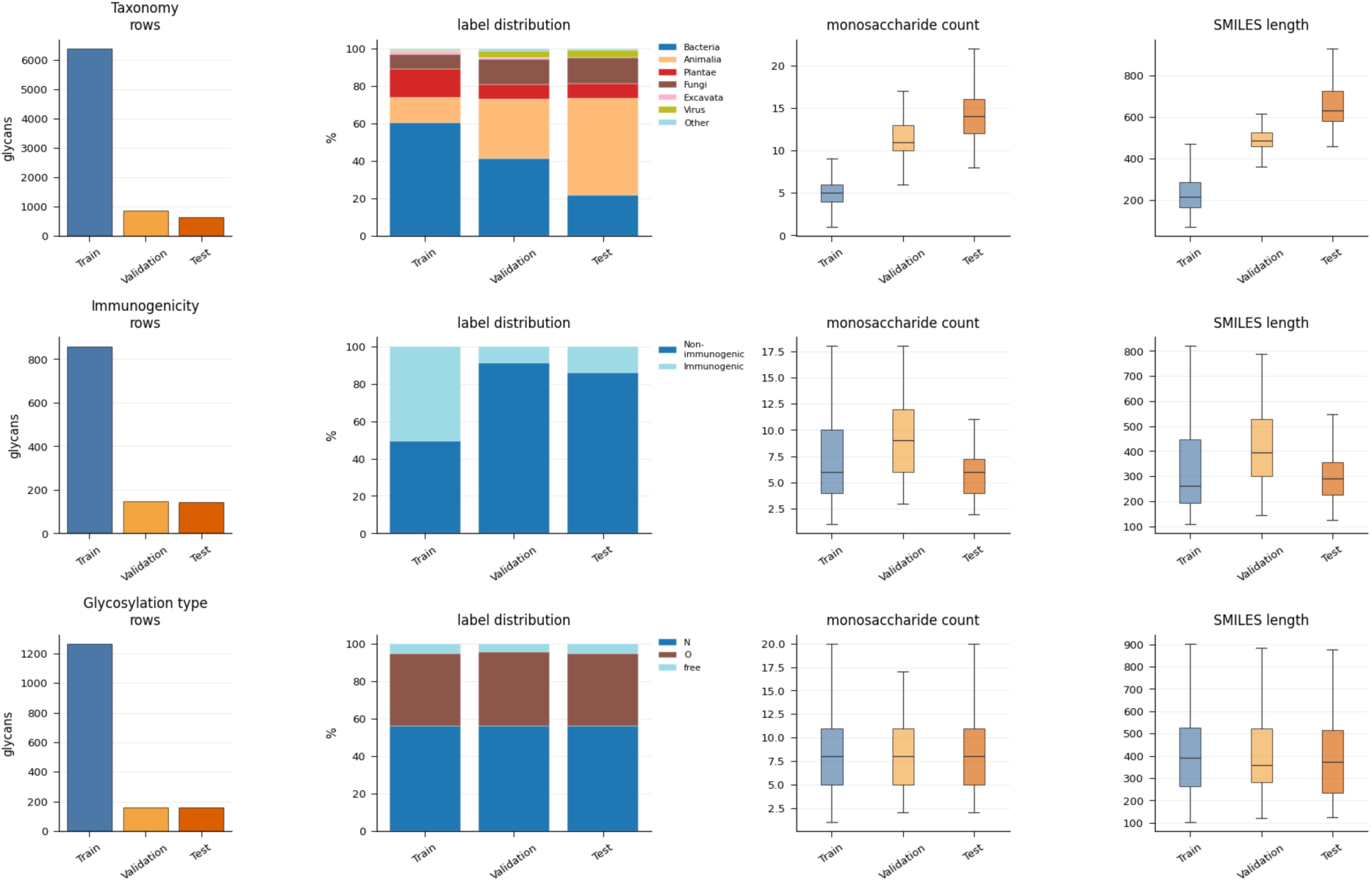
Dataset composition and class distributions across GlycanMLBenchmark tasks. Dataset characteristics for the glycan-only benchmark tasks after GlyLES conversion. For each task (taxonomy, immunogenicity and glycosylation type), panels show (left to right): the number of glycans in the training, validation and test partitions; the distribution of biological labels across each partition; the distribution of glycan size measured by monosaccharide count; and the distribution of molecular size measured by SMILES length. Taxonomy exhibits the strongest distribution shift between training and evaluation partitions, with substantially larger glycans in the validation and test sets, reflecting the benchmark’s motif-based split designed to assess structural generalization. In contrast, immunogenicity and glycosylation-type datasets show more comparable molecular-size distributions across partitions while preserving their respective class compositions.

**Supplementary Figure 2.**
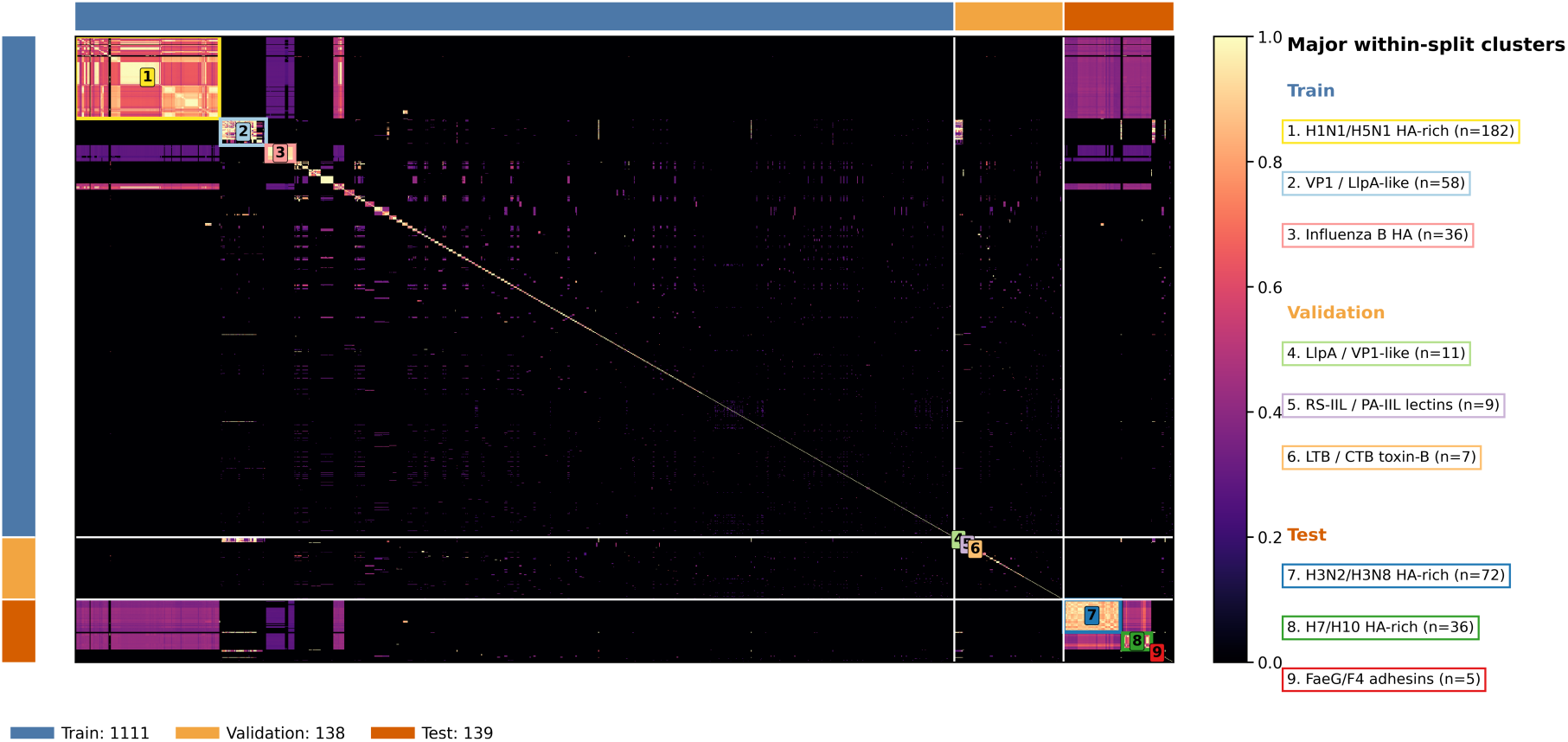
Protein sequence similarity structure underlying the protein-glycan interaction benchmark split. Pairwise protein sequence similarity matrix for the LectinOracle-derived protein-glycan interaction dataset used in the GlycanML benchmark. Proteins are ordered according to the training (blue), validation (orange) and test (red) partitions generated by MMseqs2 clustering at 50% sequence identity. Colour intensity represents normalized pairwise sequence similarity, with brighter values indicating greater sequence similarity. Major protein families, including influenza haemagglutinins, VP1 proteins, lectins and bacterial adhesins, form distinct sequence clusters. Homologous proteins are intentionally distributed across different benchmark partitions, requiring models to generalize glycan-binding specificity to evolutionarily related proteins that were not observed during training rather than memorizing individual protein-glycan interactions.

**Supplementary Figure 3.**
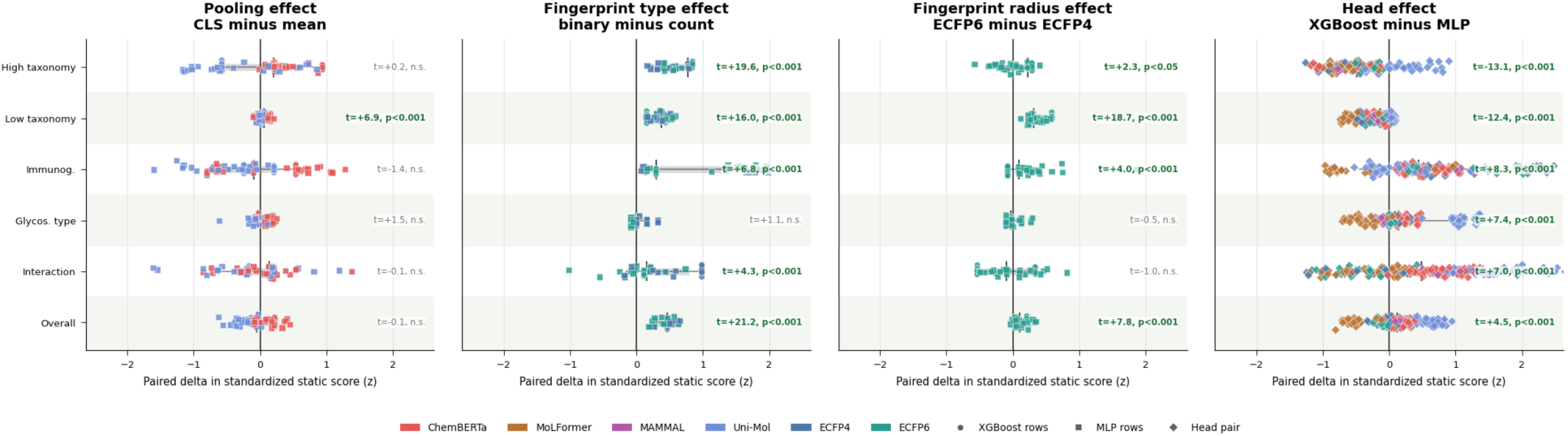
Effects of molecular representation and downstream modelling strategies on glycan prediction performance. Paired standardized performance differences for four methodological comparisons across the GlycanMLBenchmark tasks. Each point represents a matched comparison performed under identical benchmark conditions, and the x-axis reports the paired difference in standardized performance (z-score). Positive values indicate improved performance for the first method listed in each comparison. Statistical significance was assessed using paired two-sided *t*-tests across all matched model configurations. **a) Pooling strategy (CLS-mean).** Comparison of CLS-token and mean-pooled molecular embeddings for SMILES-based chemical foundation models. No consistent overall advantage was observed, although CLS pooling significantly improved low-level taxonomy prediction. **b) Comparison of binary and count-based Morgan fingerprints.** Binary fingerprints consistently outperformed count fingerprints for taxonomy, immunogenicity and protein-glycan interaction prediction, whereas glycosylation-type prediction showed no significant difference. **c) Comparison of Morgan fingerprints with radii 3 (ECFP6) and 2 (ECFP4).** Increasing fingerprint radius significantly improved taxonomy and immunogenicity prediction but provided no significant advantage for glycosylation-type or protein-glycan interaction prediction. **d) Comparison of XGBoost and multilayer perceptron (MLP) prediction heads across all static molecular representations.** XGBoost significantly improved immunogenicity, glycosylation-type and protein-glycan interaction prediction, whereas MLP performed better for both taxonomy tasks, resulting in a modest overall advantage for XGBoost. Colours indicate molecular representation families (ChemBERTa, MoLFormer, MAMMAL, Uni-Mol and Morgan fingerprints), while point shapes denote downstream prediction heads where applicable. Horizontal annotations report paired *t*-statistics and corresponding *P* values. Together, these analyses identify the methodological choices that most consistently influence downstream glycan prediction performance.

**Supplementary Figure 4.**
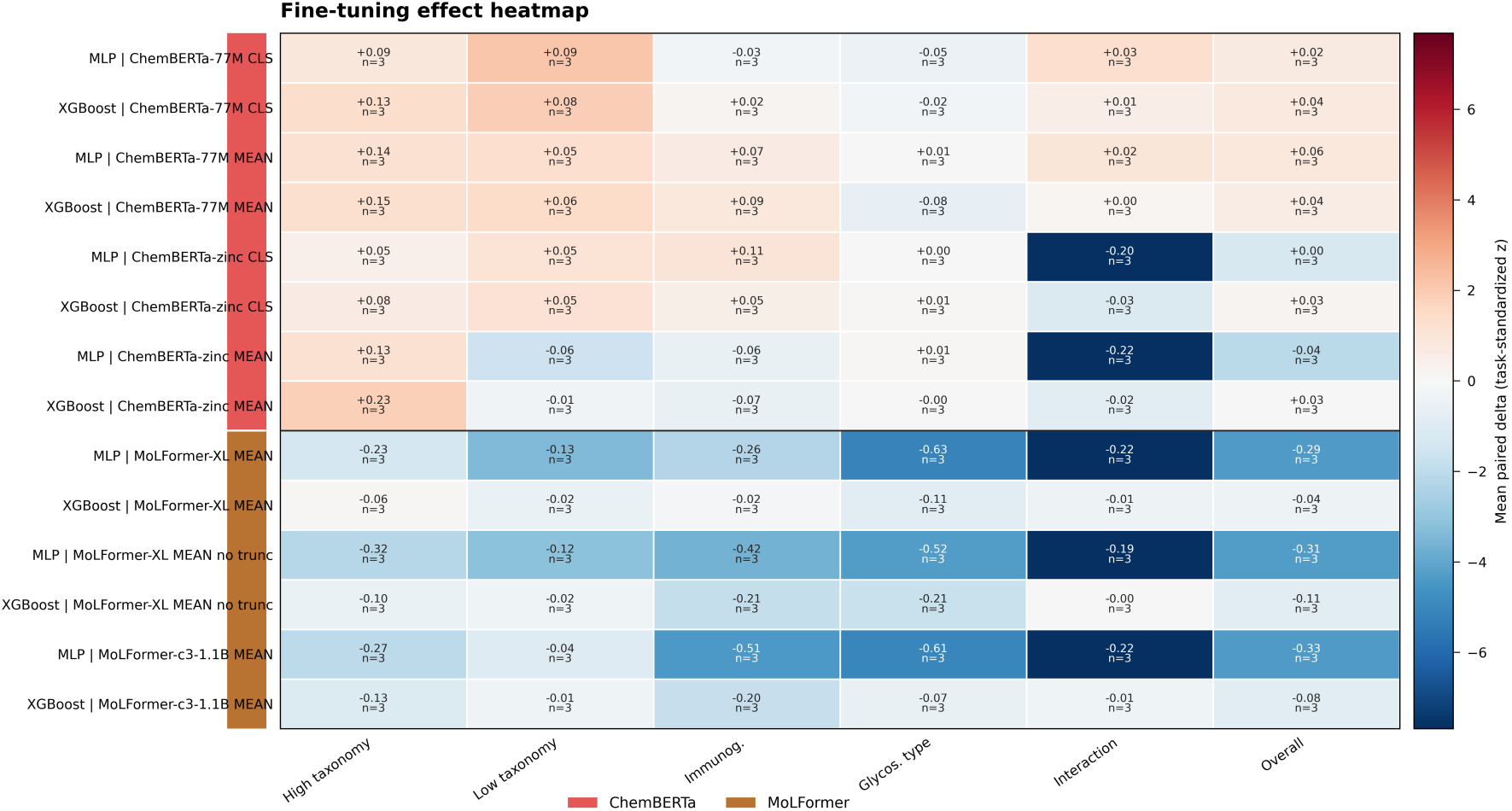
Effects of task-specific fine-tuning on chemical foundation model performance. Heatmap summarizing the effect of task-specific fine-tuning across ChemBERTa and MoLFormer model variants. Each row corresponds to a model, pooling strategy and downstream prediction head, and each column to a GlycanML benchmark task. Values represent the mean paired change in standardized performance (z-score) following fine-tuning relative to the corresponding frozen representation, averaged across three independent random seeds (*n* = 3). Positive values (red) indicate improved performance after fine-tuning, whereas negative values (blue) indicate reduced performance. Fine-tuning produced consistent performance gains for ChemBERTa models, particularly for high-level taxonomy, immunogenicity and protein-glycan interaction prediction, whereas MoLFormer models generally showed smaller or negative changes across benchmark tasks. These results demonstrate that the ability to exploit biological supervision depends strongly on the underlying molecular representation and provides the basis for the latent-space analyses presented in the main text.

**Supplementary Figure 5.**
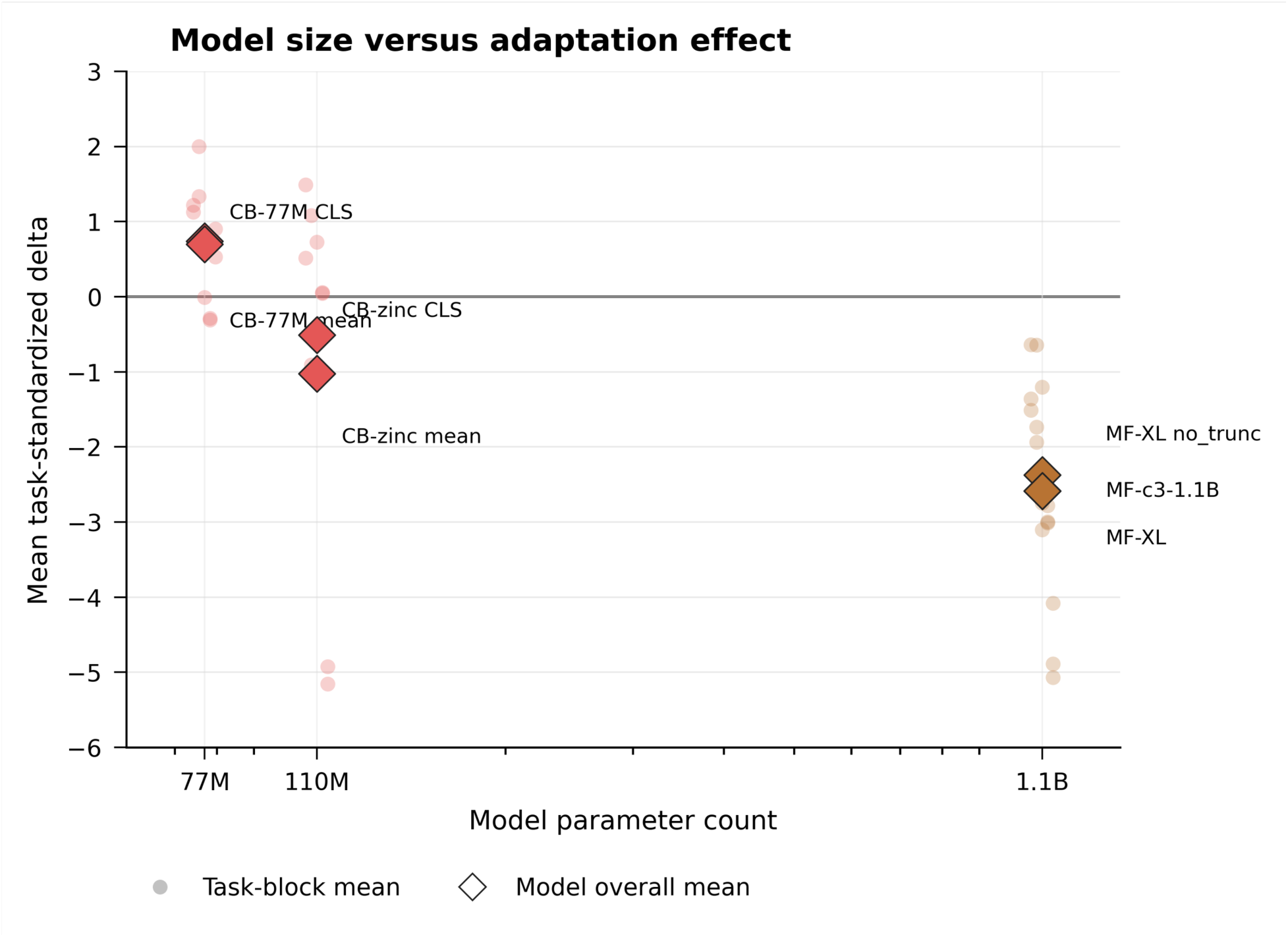
Relationship between model scale and adaptation to glycan biology. Relationship between chemical foundation model size and the effect of task-specific fine-tuning across the GlycanML benchmark tasks. The x-axis shows the approximate number of model parameters, and the y-axis reports the mean paired improvement in standardized performance (z-score) following fine-tuning relative to the corresponding frozen representation. Small circles represent individual benchmark tasks, whereas diamonds indicate the overall mean adaptation effect for each model across all tasks. ChemBERTa-77M exhibited the largest overall improvement despite having substantially fewer parameters than several larger models, whereas larger MoLFormer models generally showed limited or negative gains following fine-tuning. These results indicate that successful transfer to glycobiology depends primarily on the suitability of the learned molecular representation rather than model scale alone.

**Supplementary Figure 6.**
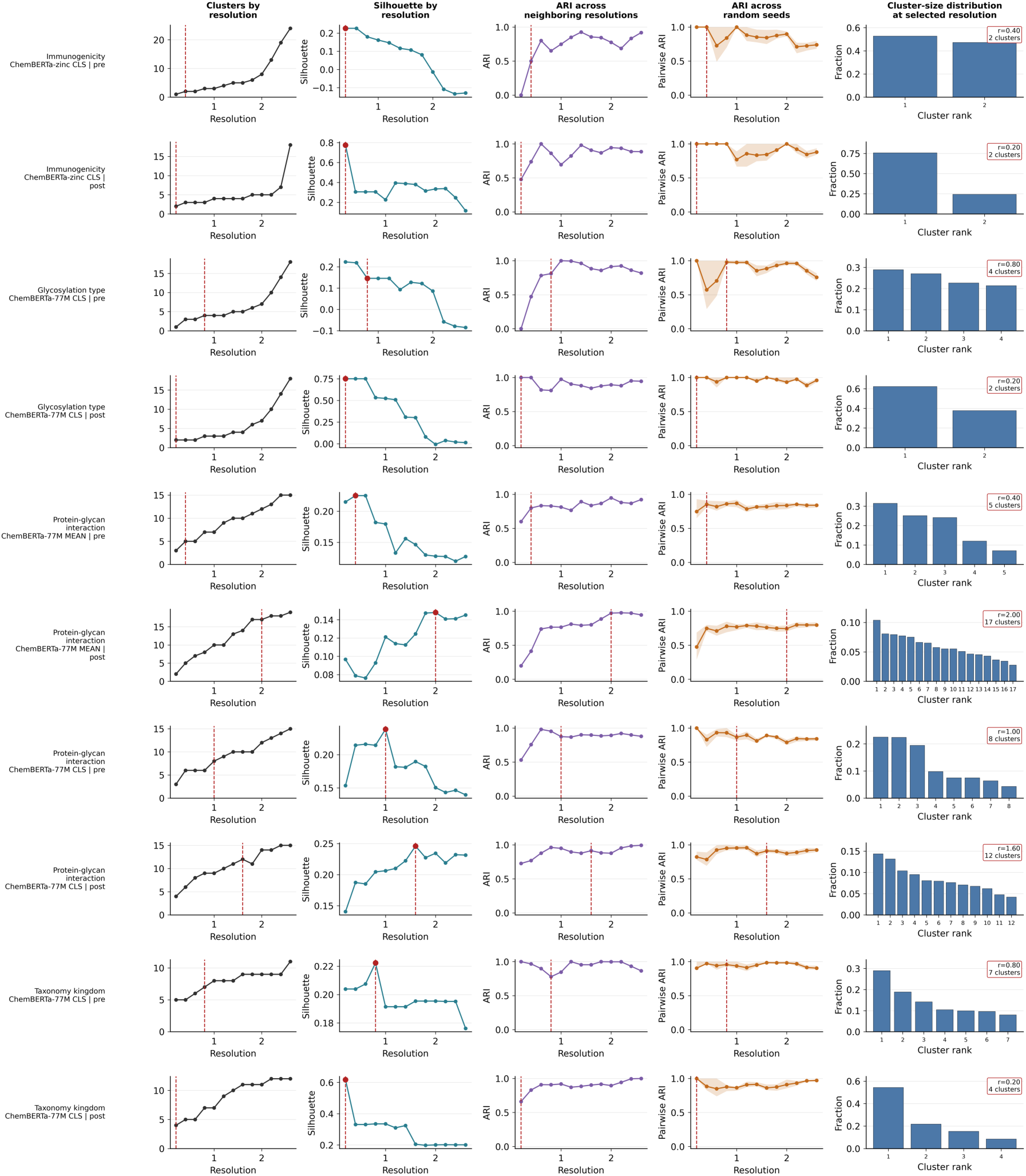
Selection of Leiden clustering resolution for embedding-space analysis. Resolution optimization for Leiden community detection across the embedding spaces analysed in this study. Each row corresponds to one pretrained or fine-tuned representation used for downstream biological interpretation. Columns show (left to right): the number of detected Leiden communities as a function of the resolution parameter; mean silhouette score calculated in the original embedding space; adjusted Rand index (ARI) between neighbouring resolutions; ARI across independent Leiden random seeds; and the cluster-size distribution obtained at the selected resolution. Red dashed lines indicate the final resolution chosen for each embedding space. Resolutions were selected using the lowest value whose silhouette score was within 0.02 of the maximum observed value, while maintaining high clustering stability across neighbouring resolutions and random seeds and avoiding excessive fragmentation into small communities. This procedure produced stable, biologically interpretable clusterings while allowing the number of communities to adapt to the intrinsic organization of each embedding space.

**Supplementary Figure 7.**
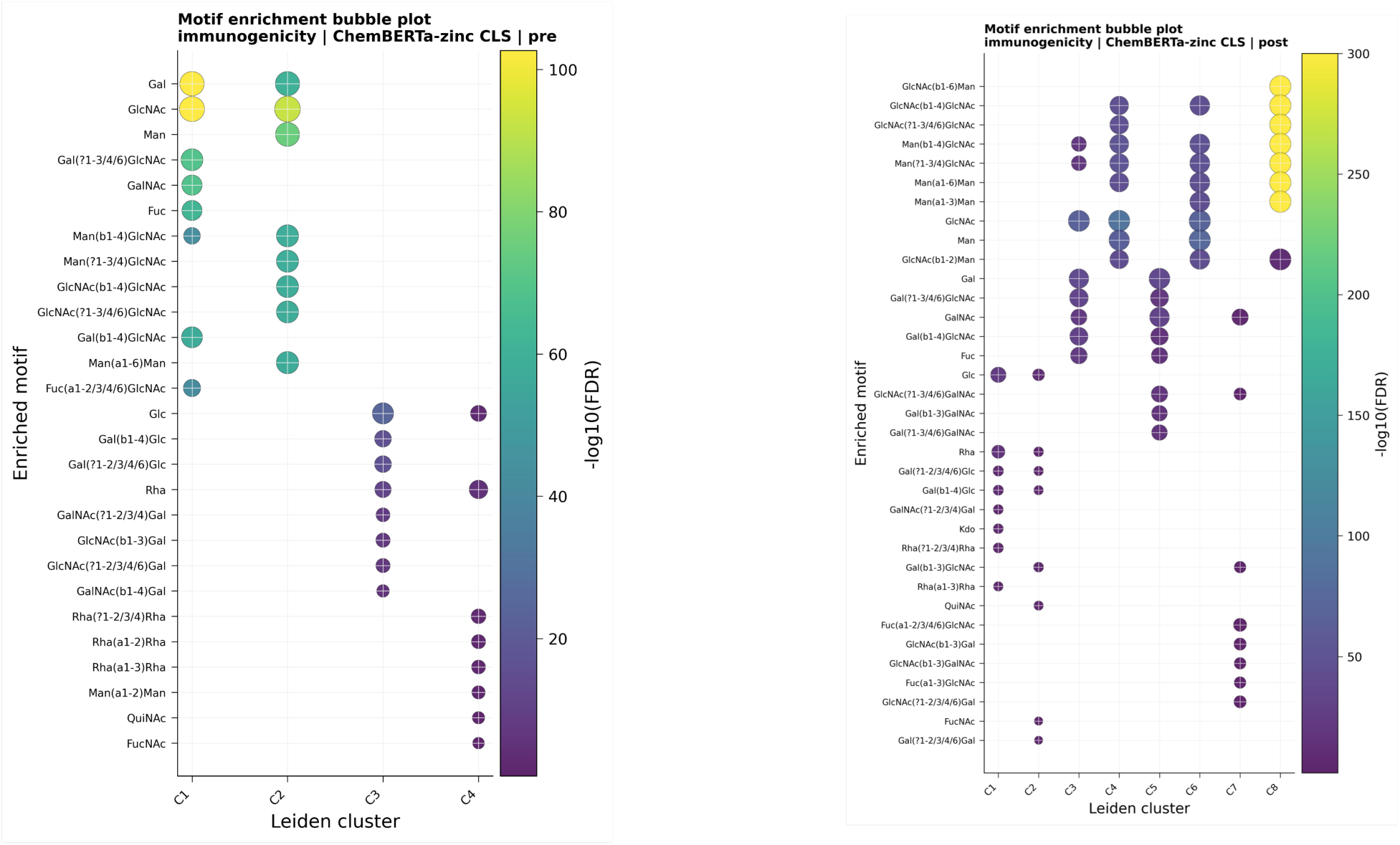
Monosaccharide and linkage enrichment of immunogenicity-associated embedding communities. Cluster-versus-rest enrichment analysis of monosaccharide identities and glycosidic linkages for the pretrained and fine-tuned ChemBERTa-zinc CLS embedding spaces used in the immunogenicity task. Each bubble represents a monosaccharide or linkage significantly enriched within a Leiden community following multiple-testing correction; colour indicates statistical significance (−log10(FDR)), and bubble size represents enrichment magnitude. Before fine-tuning, the predominantly non-immunogenic communities were enriched for Gal-, GlcNAc-, GalNAc-, Fuc- and Man-containing motifs together with canonical N-glycan and LacNAc-associated linkages, whereas the immunogenic communities were characterized by Glc-, Rha-, QuiNAc- and FucNAc-containing motifs and Gal-Glc or Rha-Rha linkages. Following fine-tuning, these broad structural signals were reorganized into more specific communities, with canonical Man/GlcNAc-rich N-glycan motifs concentrated in non-immunogenic clusters and plant-, microbial- and rare-sugar-associated motifs consolidated within the immunogenic communities. These analyses provide the monosaccharide- and linkage-level interpretation underlying the higher-order glycan motif enrichments presented in Figure 3.

**Supplementary Figure 8.**
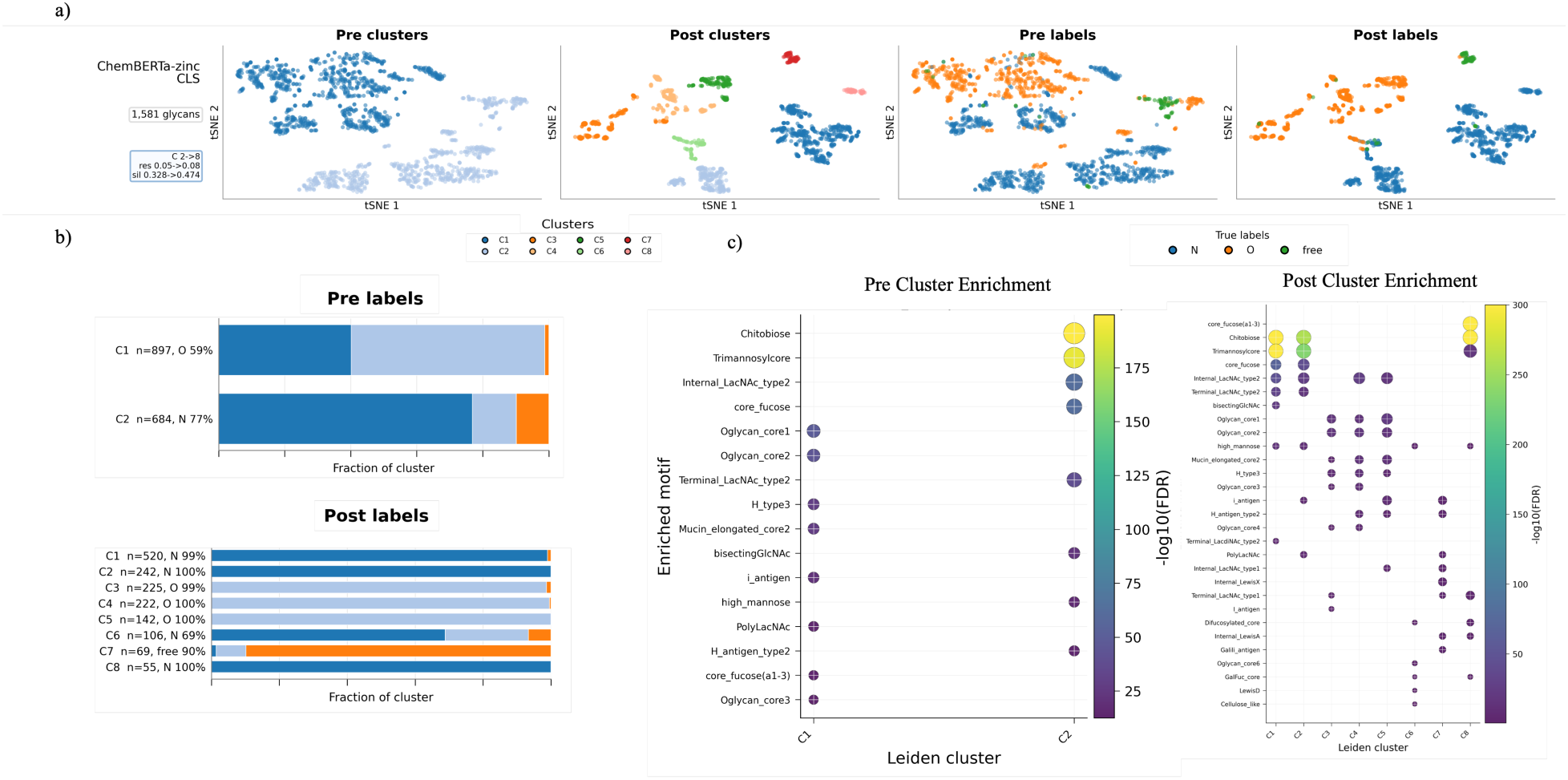
Fine-tuning reorganizes chemical representations into glycosylation-type-associated molecular communities. **a)** t-SNE projections of ChemBERTa-zinc CLS glycan embeddings before and after glycosylation-type fine-tuning. Leiden communities were identified in the original high-dimensional embedding space and projected onto the two-dimensional t-SNE visualization. Embeddings are coloured either by Leiden community or by the benchmark glycosylation-type labels (N-linked, O-linked or free glycans). Before fine-tuning, the representation broadly separated N-linked and O-linked glycans. After fine-tuning, the embedding reorganized into multiple communities corresponding to distinct N-glycan, O-glycan and free-glycan structural programmes. **b)** Glycosylation-type composition of each Leiden community before and after fine-tuning. Bars show the fraction of N-linked, O-linked and free glycans within each community, together with the dominant class and its proportion. Before fine-tuning, two broad communities captured predominantly O-linked and N-linked glycans. After fine-tuning, N-linked glycans separated into multiple canonical and branched N-glycan communities, O-linked glycans resolved into distinct mucin-type and extended O-glycan communities, and free glycans formed a dedicated community while a single mixed community retained terminal LacNAc- and sialylation-associated structures shared across attachment classes. **c)** Cluster-versus-rest enrichment of annotated glycan motifs before and after fine-tuning. Bubble plots show significantly enriched glycan motifs within each Leiden community; colour indicates statistical significance (−log10(FDR)), and bubble size represents enrichment magnitude. Fine-tuning refined the broad N-linked versus O-linked distinction into communities corresponding to canonical N-glycan cores, branched N-glycans, mucin-type O-glycans, extended O-glycans and free glycans, consistent with established glycan biosynthetic grammar.

**Supplementary Figure 9.**
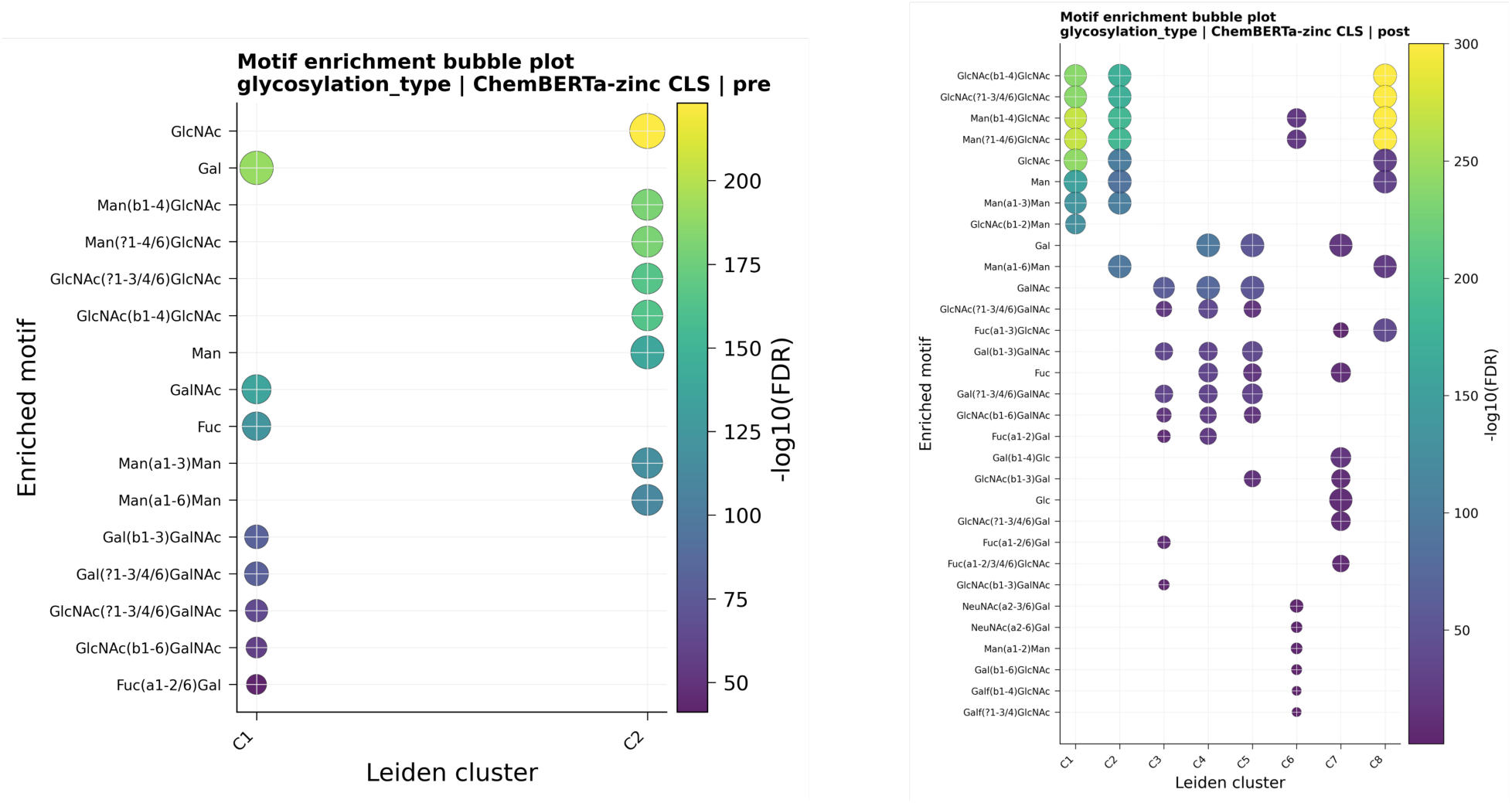
Monosaccharide and linkage enrichment within glycosylation-type-associated embedding communities. Cluster-versus-rest enrichment analysis of monosaccharide identities and glycosidic linkages for the pretrained and fine-tuned ChemBERTa-zinc CLS embedding spaces used for glycosylation-type prediction. Each bubble represents a monosaccharide or linkage significantly enriched within a Leiden community following multiple-testing correction; colour indicates statistical significance (−log10(FDR)), and bubble size represents enrichment magnitude. Before fine-tuning, the two major communities broadly separated GalNAc-centred O-glycan motifs from Man/GlcNAc-centred N-glycan motifs. Following fine-tuning, these broad structural programmes resolved into multiple communities corresponding to canonical N-glycan cores, branched and extended N-glycans, mucin-type O-glycans, terminally elaborated O-glycans and free glycans.

**Supplementary Figure 10.**
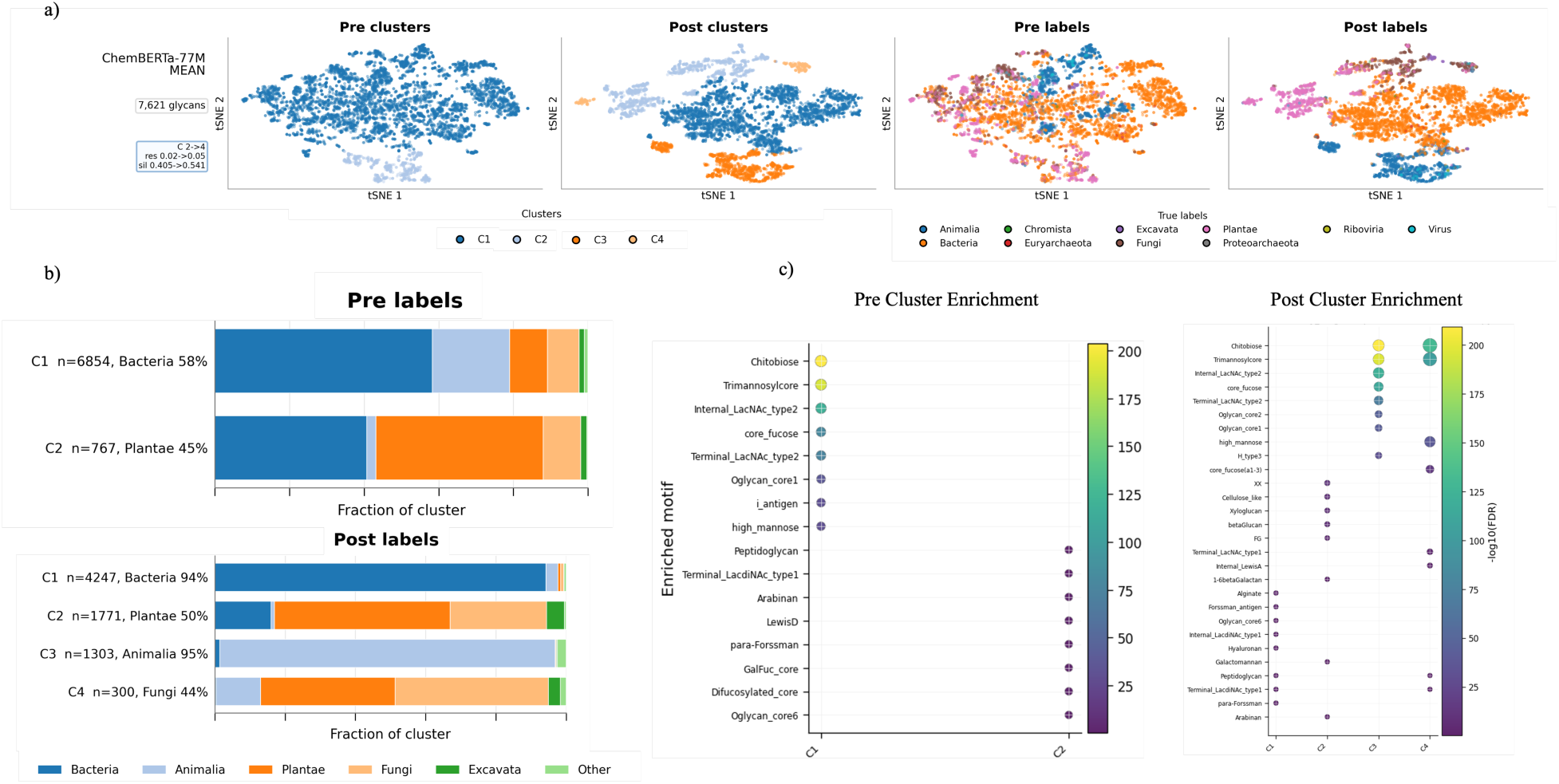
Fine-tuning reorganizes chemical representations into kingdom-associated glycan communities. **a)** t-SNE projections of ChemBERTa-77M mean glycan embeddings before and after kingdom fine-tuning. Leiden communities were identified in the original high-dimensional embedding space and projected onto the two-dimensional t-SNE visualization. Embeddings are coloured either by Leiden community or by kingdom labels. Before fine-tuning, embeddings were organized into two broad chemical communities with only partial correspondence to taxonomy. Following fine-tuning, the representation reorganized into more distinct bacterial-, animal-, plant- and fungal-associated communities. **b)** Kingdom composition of each Leiden community before and after fine-tuning. Bars show the fraction of glycans from each kingdom within each community, together with the dominant kingdom and its proportion. Before fine-tuning, communities reflected broad molecular chemotypes with mixed taxonomic composition. After fine-tuning, one community became predominantly bacterial (94%), another predominantly animal (95%), while plant-associated glycans formed a separate community and fungal glycans occupied a more heterogeneous intermediate region, consistent with their structural overlap with other non-vertebrate carbohydrate architectures. **c)** Cluster-versus-rest enrichment of annotated glycan motifs before and after fine-tuning. Bubble plots show glycan motifs significantly enriched within each Leiden community; colour indicates statistical significance (−log10(FDR)), and bubble size represents enrichment magnitude. Before fine-tuning, communities primarily distinguished canonical eukaryotic N-glycan motifs from a heterogeneous collection of non-mammalian carbohydrate features. Fine-tuning reorganized these broad chemotypes into communities enriched for bacterial cell-envelope glycans, canonical animal N- and O-glycan architectures, plant cell-wall polysaccharides and fungal high-mannose-associated structures, providing a biologically coherent organization of kingdom-associated glycan ecologies.

**Supplementary Figure 11.**
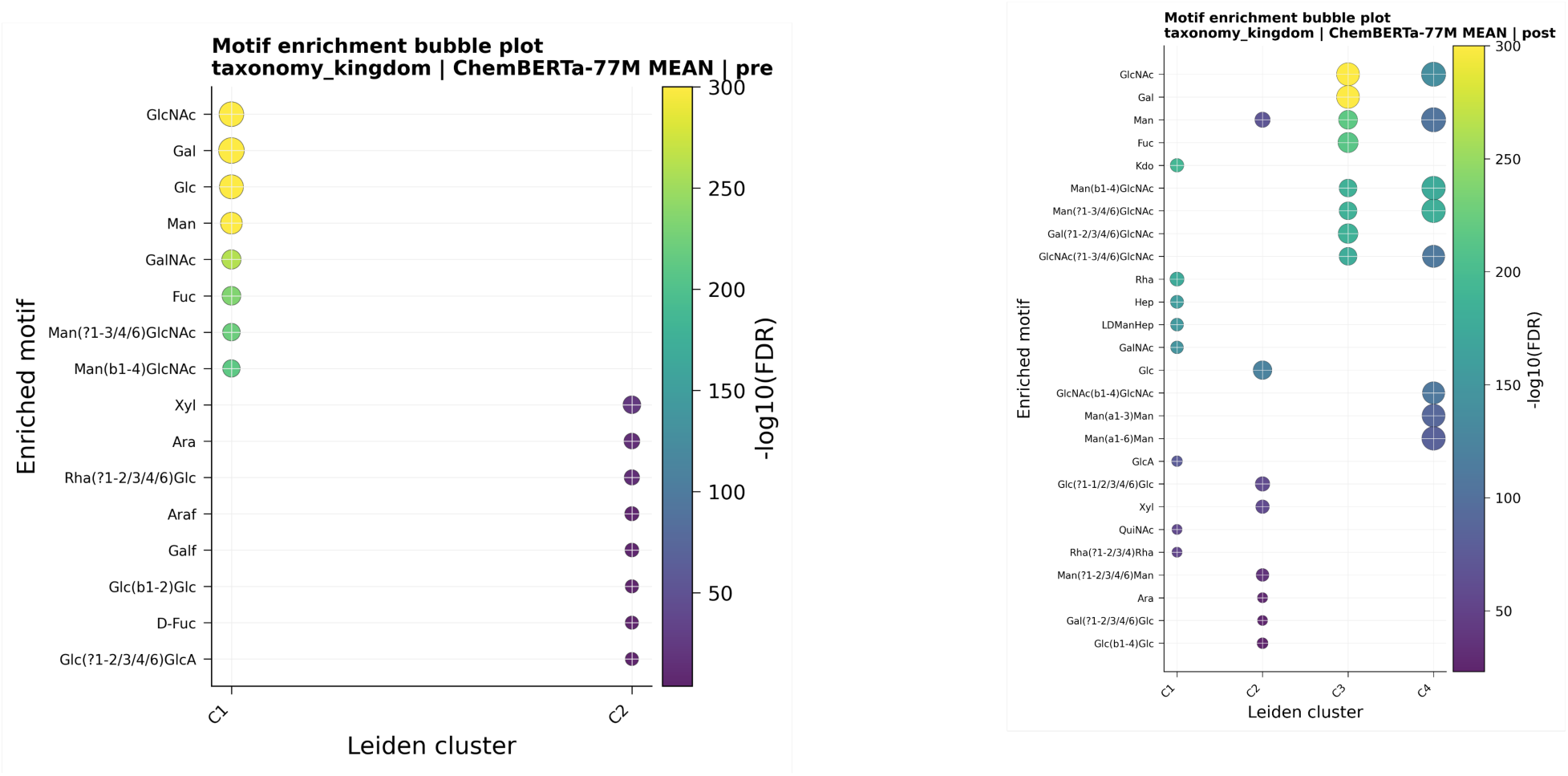
Monosaccharide and linkage enrichment of kingdom-associated embedding communities. Cluster-versus-rest enrichment analysis of monosaccharide identities and glycosidic linkages for the pretrained and fine-tuned ChemBERTa-77M mean embedding spaces used for kingdom prediction. Each bubble represents a monosaccharide or linkage significantly enriched within a Leiden community following multiple-testing correction; colour indicates statistical significance (−log10(FDR)), and bubble size represents enrichment magnitude. Before fine-tuning, the two major communities broadly distinguished common eukaryotic glycan chemistry, characterized by GlcNAc-, Gal-, Man-, GalNAc- and Fuc-containing motifs together with canonical Man-GlcNAc linkages, from a heterogeneous non-mammalian carbohydrate space enriched for Xyl-, Ara-, Araf-, Galf- and rhamnose-containing motifs. Following fine-tuning, these broad structural programmes reorganized into more taxonomically coherent communities corresponding to bacterial cell-envelope glycans enriched for Kdo, Rha, Hep and L,D-heptose motifs, canonical animal N- and O-glycan architectures, plant-associated polysaccharides enriched for Glc-, Xyl- and Ara-containing structures, and a Man-rich fungal-associated community.

**Supplementary Figure 12.**
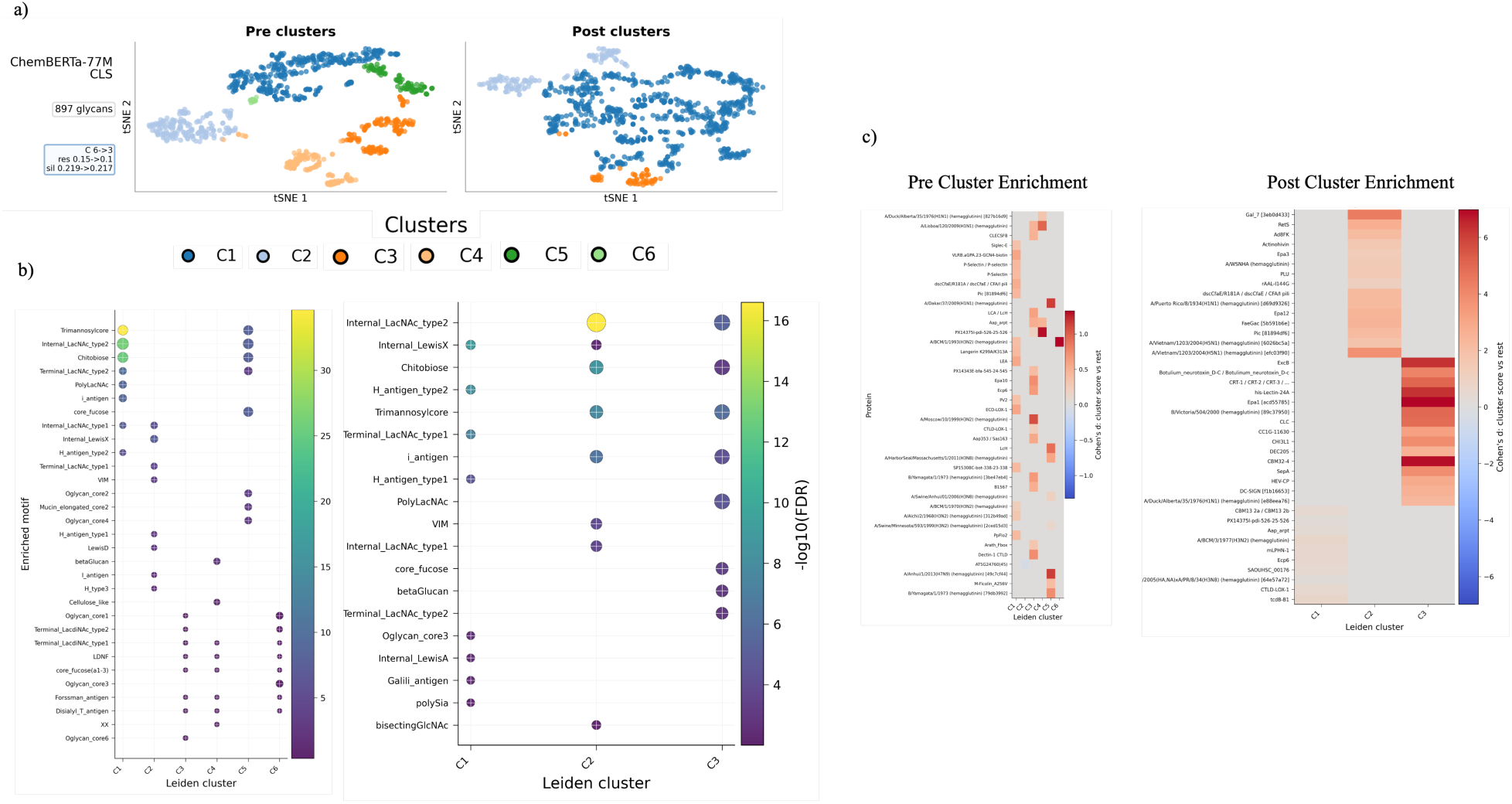
Fine-tuning reorganizes chemical representations into protein-recognition programmes. **a)** t-SNE projections of ChemBERTa-77M CLS glycan embeddings before and after protein-glycan interaction fine-tuning. Leiden communities were identified in the original high-dimensional embedding space and projected onto the two-dimensional t-SNE visualization. Unlike the classification tasks, no global interaction labels are shown because the benchmark labels are defined for glycan-protein pairs rather than for glycans alone. Fine-tuning reorganized the pretrained chemical communities into three major glycan recognition programmes. **b)** Cluster-versus-rest enrichment of annotated glycan motifs before and after fine-tuning. Bubble plots show glycan motifs significantly enriched within each Leiden community; colour indicates statistical significance (−log10(FDR)), and bubble size represents enrichment magnitude. Before fine-tuning, structural motifs were distributed across six chemically defined communities. Following fine-tuning, these reorganized into three recognition-associated programmes corresponding to terminal fucosylated/Lewis-type epitopes, sialylated Gal/LacNAc-rich receptor glycans and core N-glycan/scaffold-associated structures. Corresponding monosaccharide- and linkage-level enrichments are shown in Supplementary Fig. 13. **c)** Cluster-versus-rest protein enrichment before and after fine-tuning. Heatmaps show proteins whose interaction scores are significantly enriched for glycans within each Leiden community relative to all remaining communities (Cohen’s *d*). Before fine-tuning, associations with lectins, microbial adhesins and viral glycoproteins were distributed across multiple chemically related communities. Following fine-tuning, these associations became concentrated into three coherent recognition programmes, linking terminal fucosylated glycans, sialylated Gal/LacNAc receptor glycans and N-glycan scaffold structures to distinct groups of glycan-binding proteins. These results demonstrate that task supervision reorganizes glycan representation space into biologically interpretable protein-recognition programmes rather than simply increasing overall interaction prediction performance.

**Supplementary Figure 13.**
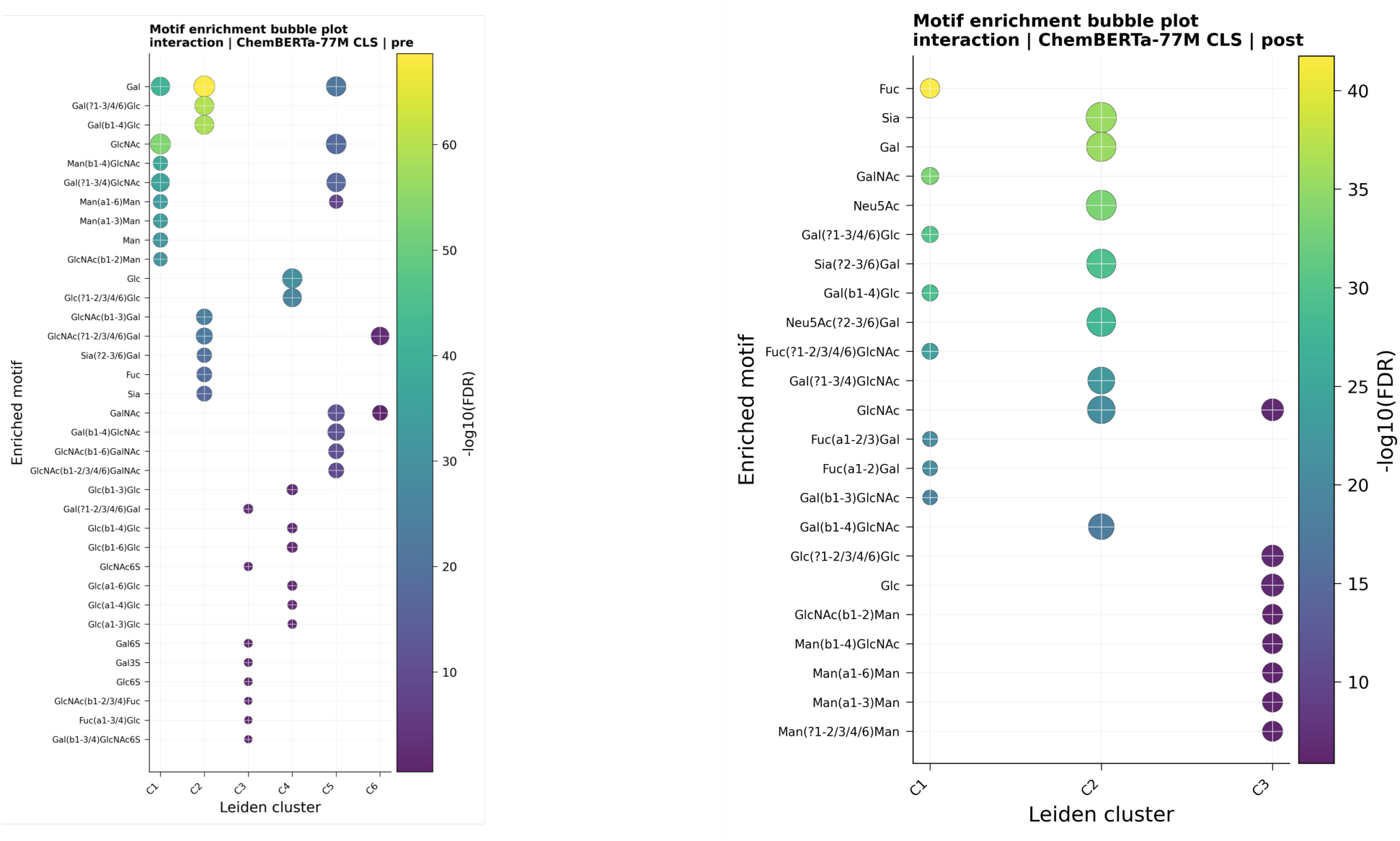
Monosaccharide and linkage enrichment of protein-recognition-associated embedding communities. Cluster-versus-rest enrichment analysis of monosaccharide identities and glycosidic linkages for the pretrained and fine-tuned ChemBERTa-77M CLS embedding spaces used for protein-glycan interaction prediction. Each bubble represents a monosaccharide or linkage significantly enriched within a Leiden community following multiple-testing correction; colour indicates statistical significance (−log10(FDR)), and bubble size represents enrichment magnitude. Before fine-tuning, terminal fucosylated, sialylated, Gal/LacNAc-rich and N-glycan scaffold motifs were distributed across multiple chemically defined communities. Following fine-tuning, these broad structural signals were reorganized into three distinct recognition programmes: a terminal fucosylated/Lewis-associated community, a sialylated Gal/LacNAc-rich receptor-glycan community enriched for Neu5Ac and α2-3/6-linked sialic acid motifs, and a core/scaffold-recognition community enriched for GlcNAc-, Man- and chitobiose-containing N-glycan motifs.

**Supplementary Figure 14.**
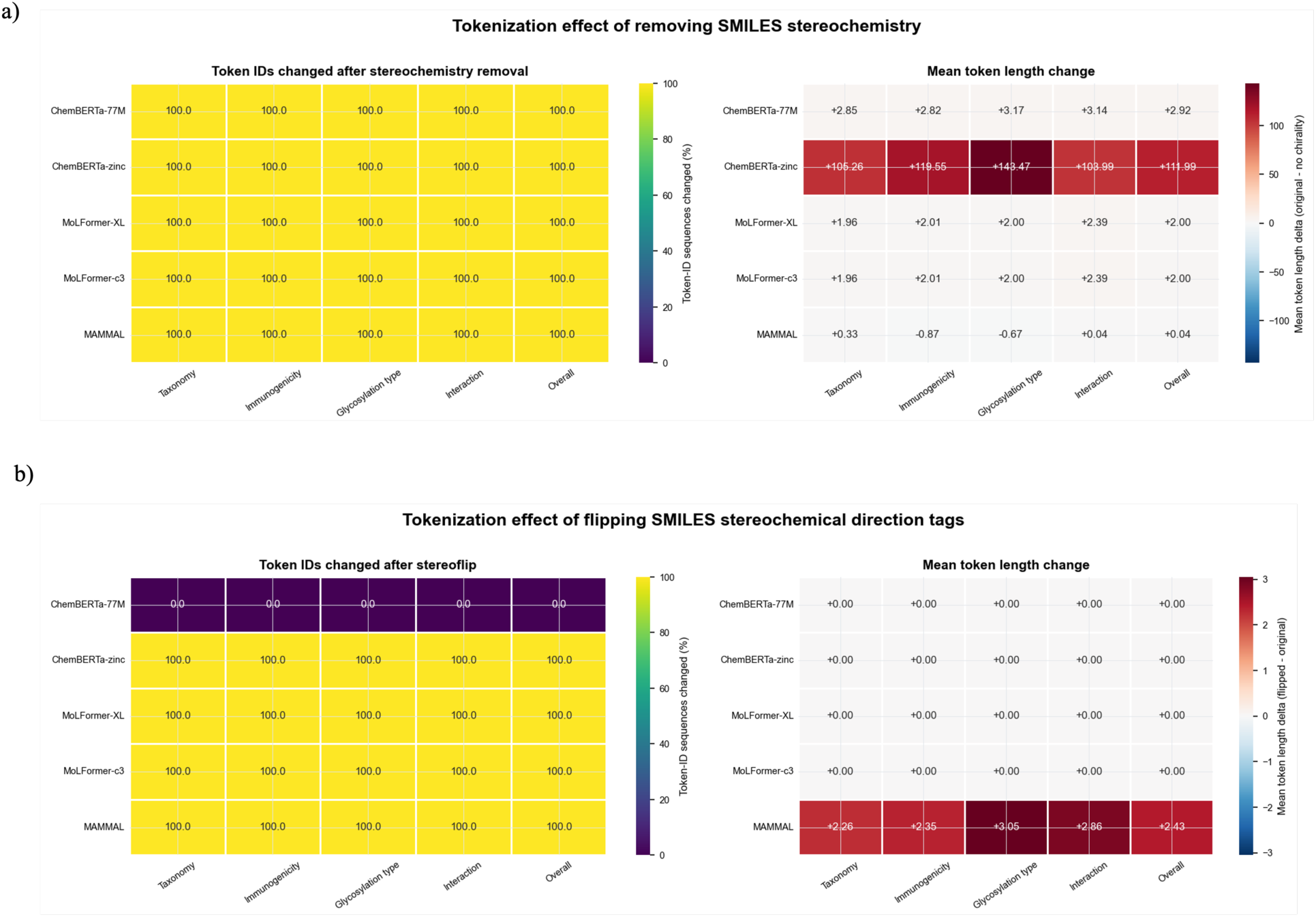
Tokenizer responses to stereochemical edits in glycan SMILES. **a)** removing stereochemical annotations changed token IDs in all models and produced the largest token-length shifts in ChemBERTa-zinc. **b)** stereochemistry inversion left ChemBERTa-77M tokenization unchanged, whereas ChemBERTa-zinc, MoLFormer and MAMMAL produced different token sequences. Left panels show the percentage of token-ID sequences changed; right panels show mean token-length differences.

**Supplementary Figure 15.**
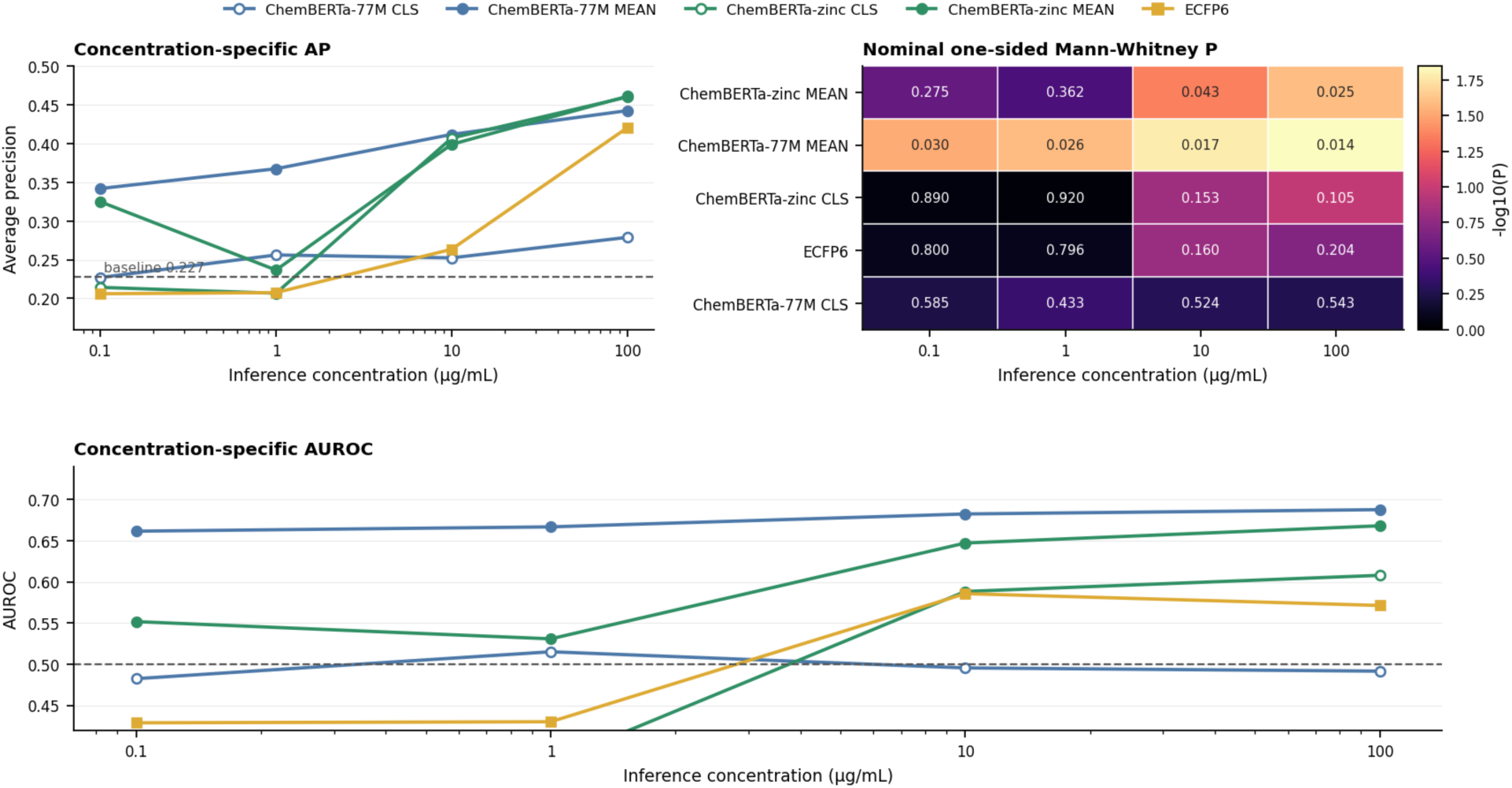
Protein Concentration dependence of LiGA binder prioritization across molecular representations. Model predictions were evaluated separately at each concentration input (0.1, 1, 10 and 100 µg/mL) using the independent LiGA panel of 66 lectin-glycan interactions, including 15 experimentally positive and 51 non-positive pairs. Top left, concentration-specific average precision (AP) for each representation. The dashed horizontal line indicates the random AP expectation of 15/66=0.227. Top right, nominal one-sided Mann-Whitney U tests assessing whether experimentally positive interactions received higher prediction scores than non-positive interactions at each protein concentration. Cell annotations report the raw nominal P-values, whereas colour intensity represents −log10(P). Bottom, concentration-specific AUROC values; the dashed horizontal line indicates chance-level discrimination (AUROC = 0.5). Filled circles denote mean-pooled ChemBERTa embeddings, open circles denote CLS representations and squares denote ECFP6 fingerprints. Mean-pooled ChemBERTa representations showed the strongest and most consistent binder-ranking signal: ChemBERTa-77M MEAN achieved the highest AUROC across concentration inputs, whereas ChemBERTa-zinc MEAN showed progressively stronger AP and positive-versus-non-positive score separation at higher concentrations.

**Supplementary Figure 16.**
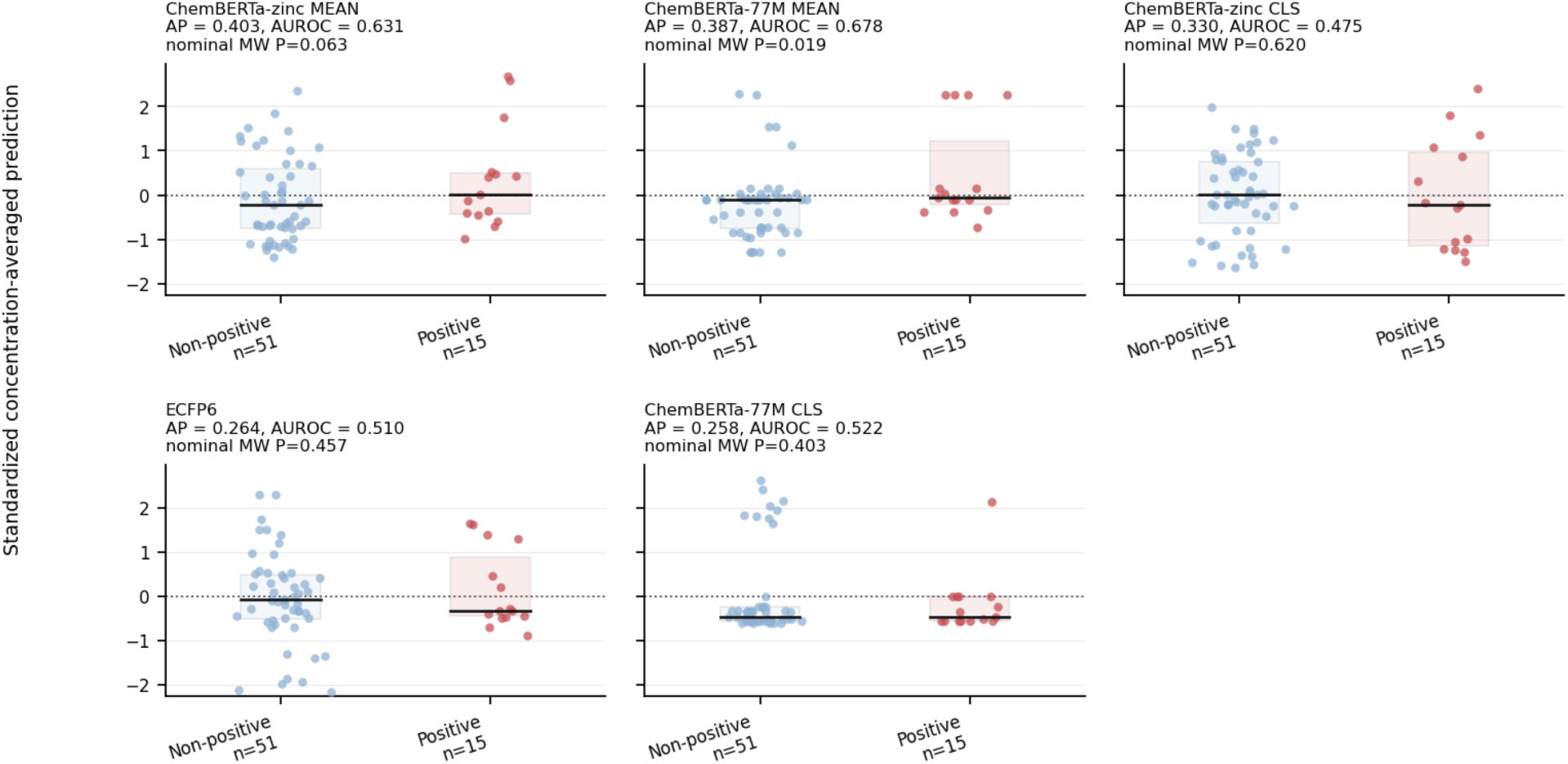
Concentration-averaged LiGA prediction-score distributions across molecular representations. Predictions for each of the 66 LiGA lectin-glycan pairs were averaged across the four concentration inputs (0.1, 1, 10 and 100 µg/mL) and standardized separately within each representation. Distributions are shown for experimentally non-positive (n=51, blue) and positive (n=15, red) interactions. Points represent individual lectin-glycan pairs, shaded boxes indicate the interquartile range, black horizontal lines indicate medians and the dotted horizontal line marks a standardized score of zero. Average precision (AP), area under the receiver-operating-characteristic curve (AUROC) and nominal one-sided Mann-Whitney P-values testing whether positive interactions received higher scores than non-positive interactions are reported above each panel. Mean-pooled ChemBERTa representations showed the strongest binder-prioritization signal: ChemBERTa-zinc MEAN achieved the highest AP (0.403), whereas ChemBERTa-77M MEAN achieved the highest AUROC (0.678) and strongest nominal positive-versus-non-positive score separation (P=0.019). CLS representations and ECFP6 showed greater overlap between the two experimental classes.

### Supplementary Tables

**Supplementary Table 1.**
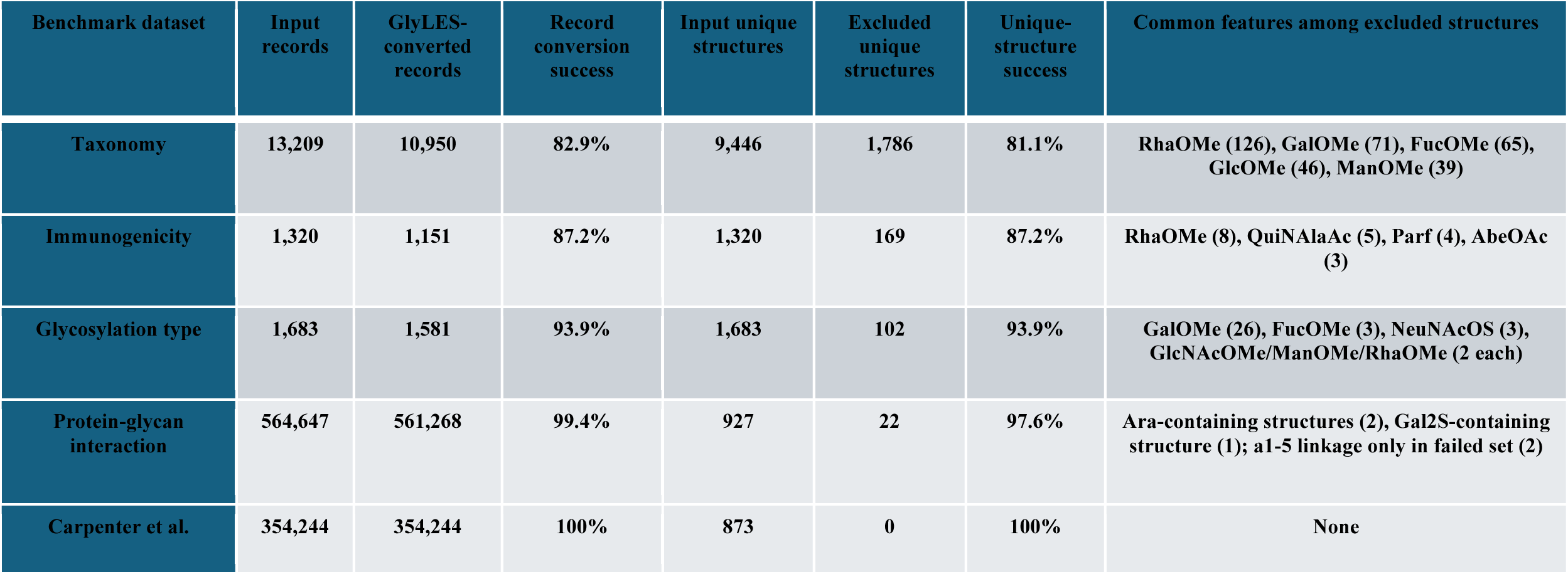
GlyLES conversion statistics across Datasets. Summary of GlyLES conversion performance across all datasets used in this study. For each dataset, the table reports the total number of input records, the number of successfully converted records, record-level conversion success, the number of unique glycan structures before and after conversion, unique-structure conversion success, and the most frequent structural features associated with conversion failures. The protein-glycan interaction benchmark showed the highest conversion success among the benchmark datasets (99.4% of records and 97.6% of unique structures), whereas taxonomy displayed the greatest structural diversity and, accordingly, the largest number of conversion failures. In contrast, the Carpenter et al. dataset was converted in full, with 100% success at both the record and unique-structure levels. Across datasets with failed conversions, the most common problematic features included uncommon modified monosaccharides, methylated sugars, and unsupported linkage patterns, indicating that conversion failures were mainly associated with rare glycan chemistries rather than canonical mammalian glycan structures.

**Supplementary Table 2.**
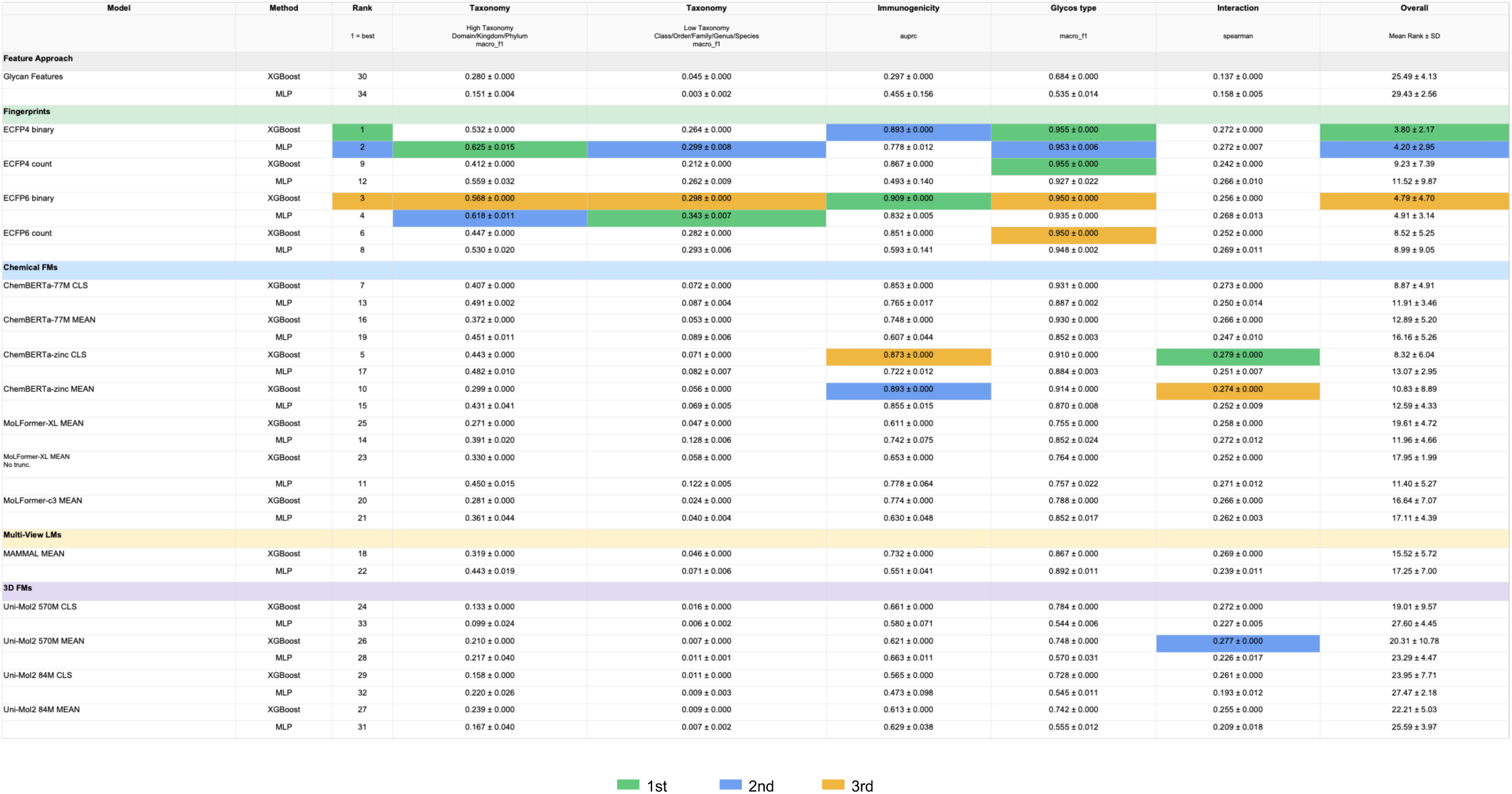
Performance of molecular representations across the GlycanML benchmark - Static Setting. Performance of all molecular representations evaluated on the GlycanML benchmark taxonomy, immunogenicity, glycosylation-type and protein-glycan interaction tasks for the static setting. Results are shown separately for static molecular representations, including engineered glycan graph features, frozen chemical foundation model (ChemFM) embeddings, Morgan fingerprints, multi-view molecular models and three-dimensional molecular foundation models. Taxonomy performance is reported as macro-F1 for each taxonomic level, immunogenicity as area under the precision-recall curve (AUPRC), glycosylation type as macro-F1, and protein-glycan interaction prediction as Spearman’s correlation. Values represent mean ± standard deviation. For each benchmark task, the highest, second-highest and third-highest performing models are highlighted in green, blue and orange, respectively.

**Supplementary Table 3.**
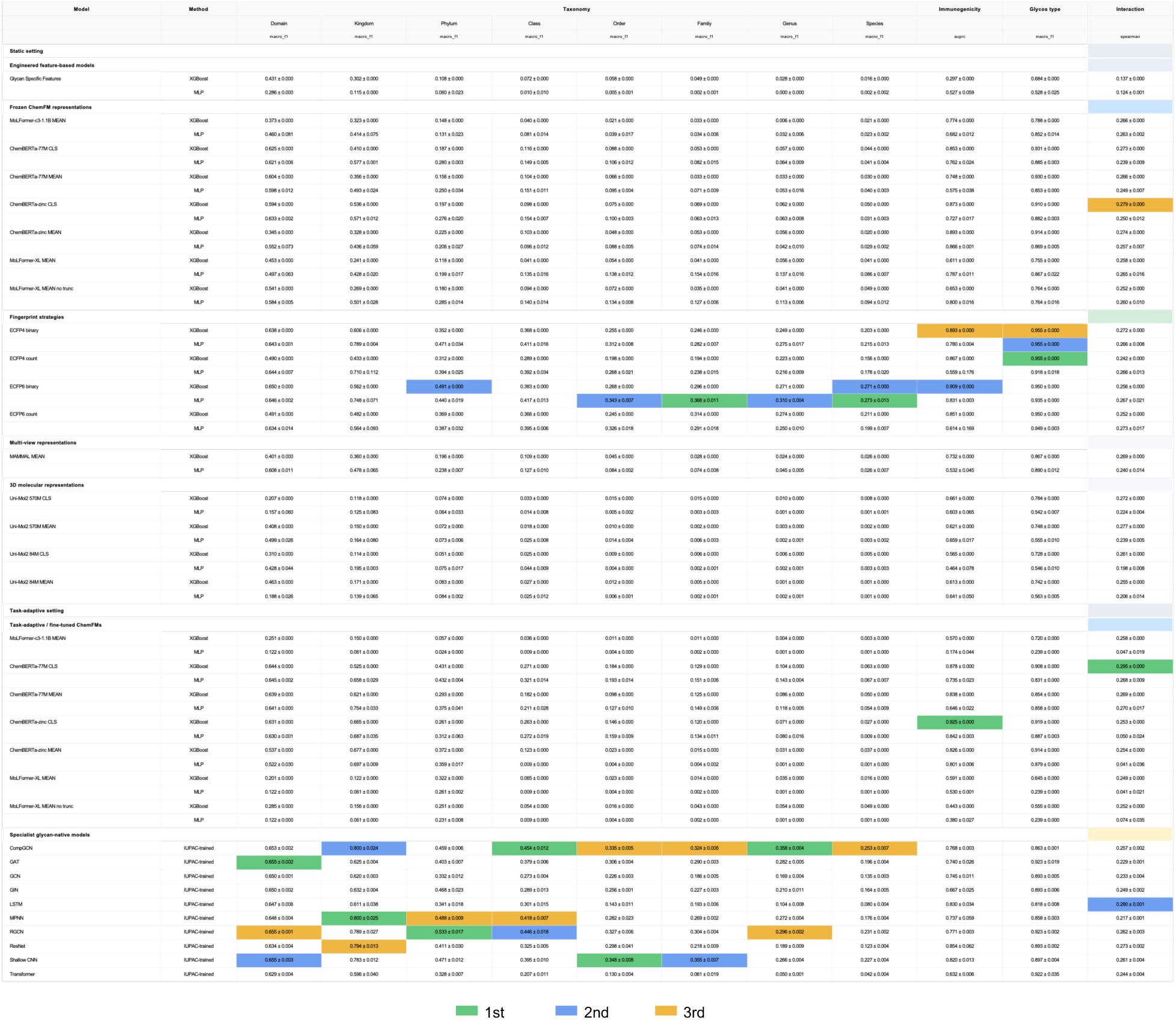
Performance of molecular representations across the GlycanML benchmark - Static and Full Setting. Performance of all molecular representations evaluated on the GlycanML benchmark taxonomy, immunogenicity, glycosylation-type and protein-glycan interaction tasks. Results are shown separately for static molecular representations, including engineered glycan graph features, frozen chemical foundation model (ChemFM) embeddings, Morgan fingerprints, multi-view molecular models and three-dimensional molecular foundation models, and task-adaptive models, including fine-tuned chemical foundation models and glycan-native models trained on IUPAC-Condensed sequences or glycan graphs. Taxonomy performance is reported as macro-F1 for each taxonomic level, immunogenicity as area under the precision-recall curve (AUPRC), glycosylation type as macro-F1, and protein-glycan interaction prediction as Spearman’s correlation. Values represent mean ± standard deviation. For each benchmark task, the highest, second-highest and third-highest performing models are highlighted in green, blue and orange, respectively. This table provides a complete overview of model performance across all benchmark tasks and molecular representation strategies.

**Supplementary Table 4.** Characteristics of the pretrained foundation models evaluated in this study. Summary of the molecular and protein foundation models included in the benchmark. For each model, the table reports the pretrained checkpoint, parameter count, backbone architecture, tokenizer or molecular featurizer, and the effective context length used during embedding extraction. For SMILES-based chemical foundation models, the reported context length corresponds to the maximum sequence length used during inference, whereas for Uni-Mol it reflects the maximum number of non-hydrogen atoms processed by the atom-level encoder. These implementation details define the effective molecular context available to each model and provide the basis for the tokenizer, truncation and representation analyses presented in the main text.

| Foundational Model | Checkpoint used | Parameters (M) | Architecture / size | Tokenizer or featurizer | Context length used |
| --- | --- | --- | --- | --- | --- |
| ChemBERTa-zinc-base-v1 | seyonec/ChemBERTa-zinc-base-v1 | 44 | RoBERTa MLM; 6 layers; hidden size 768 | RoBERTa byte-level BPE; tokenizer length 767 | 512 SMILES tokens; truncation enabled |
| ChemBERTa-77M-MLM | DeepChem/ChemBERTa-77M-MLM | 3.43 | RoBERTa MLM; 3 layers; hidden size 384 | RoBERTa byte-level BPE; tokenizer length 593; model vocab size 600 | 512 SMILES tokens; truncation enabled |
| MoLFormer-XL-both-10pct | ibm/MoLFormer-XL-both-10pct | 46.8 | MoLFormer MLM; 12 layers; hidden size 768 | MoLformerTokenizer; SMILES WordLevel tokenizer; vocab size 2,362 | 202 SMILES tokens for truncated runs; extra mean no-truncation variant also extracted |
| MoLFormer-c3-1.1B | DeepChem/MoLFormer-c3-1.1B | 46.8 | MoLFormer MLM; 12 layers; hidden size 768 | MoLformerTokenizer; SMILES WordLevel tokenizer; vocab size 2,362 | 202 SMILES tokens; truncation enabled |
| MAMMAL | ibm/biomed.omics.bl.sm.ma-ted-458m | 458 | T5-style multimodal encoder; d_model 768; 12 layers; encoder head 2048/1024 | ModularTokenizerOp; SMILES BPE component plus AA, gene and cell-attribute tokenizers | No explicit max_seq_len in static embedding extraction; no truncation recorded |
| Uni-Mol2 570M | UniMol/weights/modelzoo/570M/checkpoint.pt | 570 | Uni-Mol2; 32 layers; hidden size 1,536 | UniMolV2Feature atom/graph featurizer; hydrogens removed | 128 no-H atoms; structures above max_atoms are cropped by UniMolV2Feature |
| Uni-Mol2 84M | UniMol/weights/modelzoo/84M/checkpoint.pt | 84 | Uni-Mol2; 12 layers; hidden size 768 | UniMolV2Feature atom/graph featurizer; hydrogens removed | 128 no-H atoms; structures above max_atoms are cropped by UniMolV2Feature |
| ESM2 | facebook/esm2_t33_650M_UR50D | 652 | ESM-2 masked language model; 33 layers; hidden size 1,280 | EsmTokenizer amino-acid tokenizer; vocab size 33 | 1,026 protein tokens; truncation enabled |

**Supplementary Table 5.**
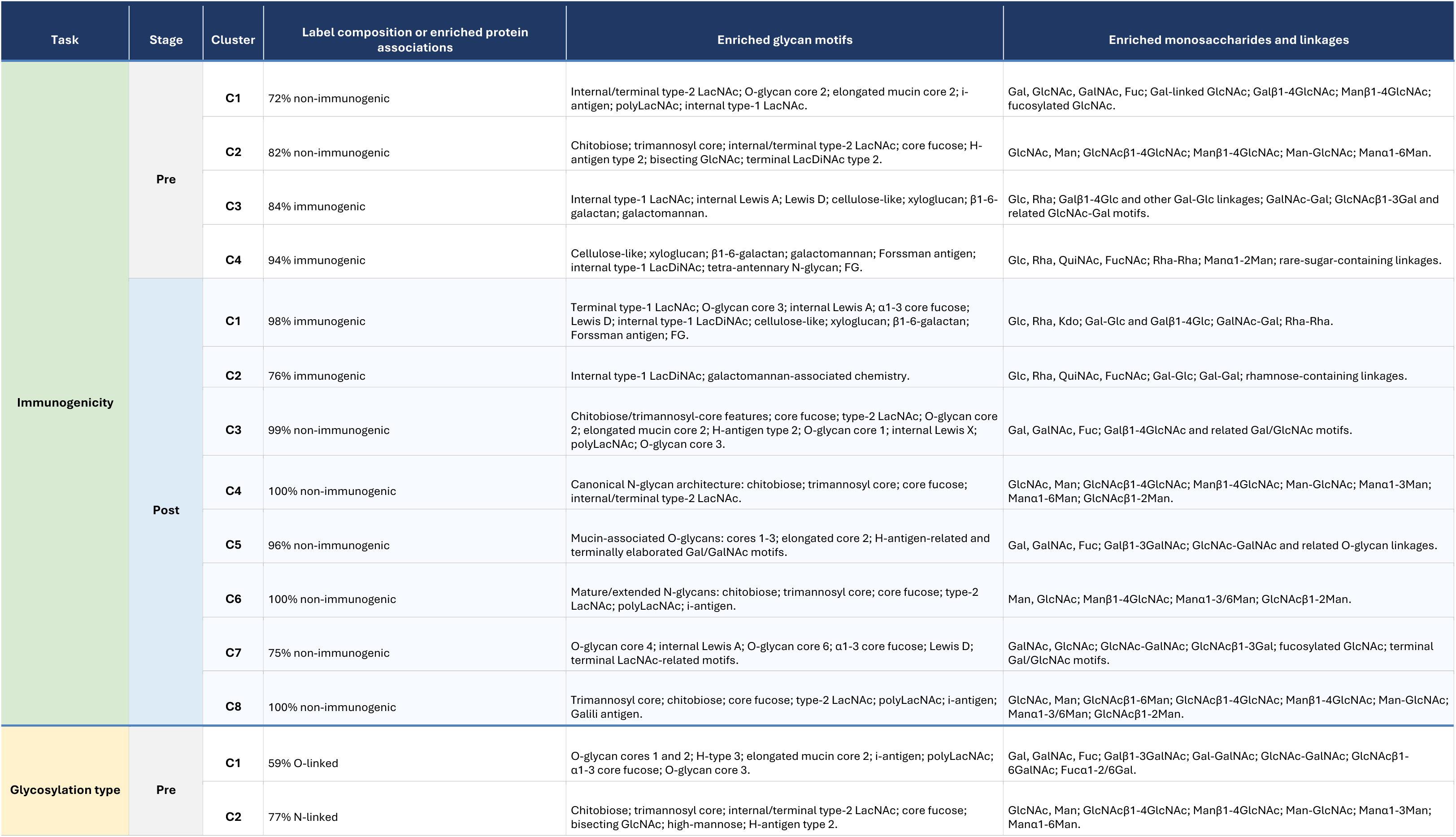
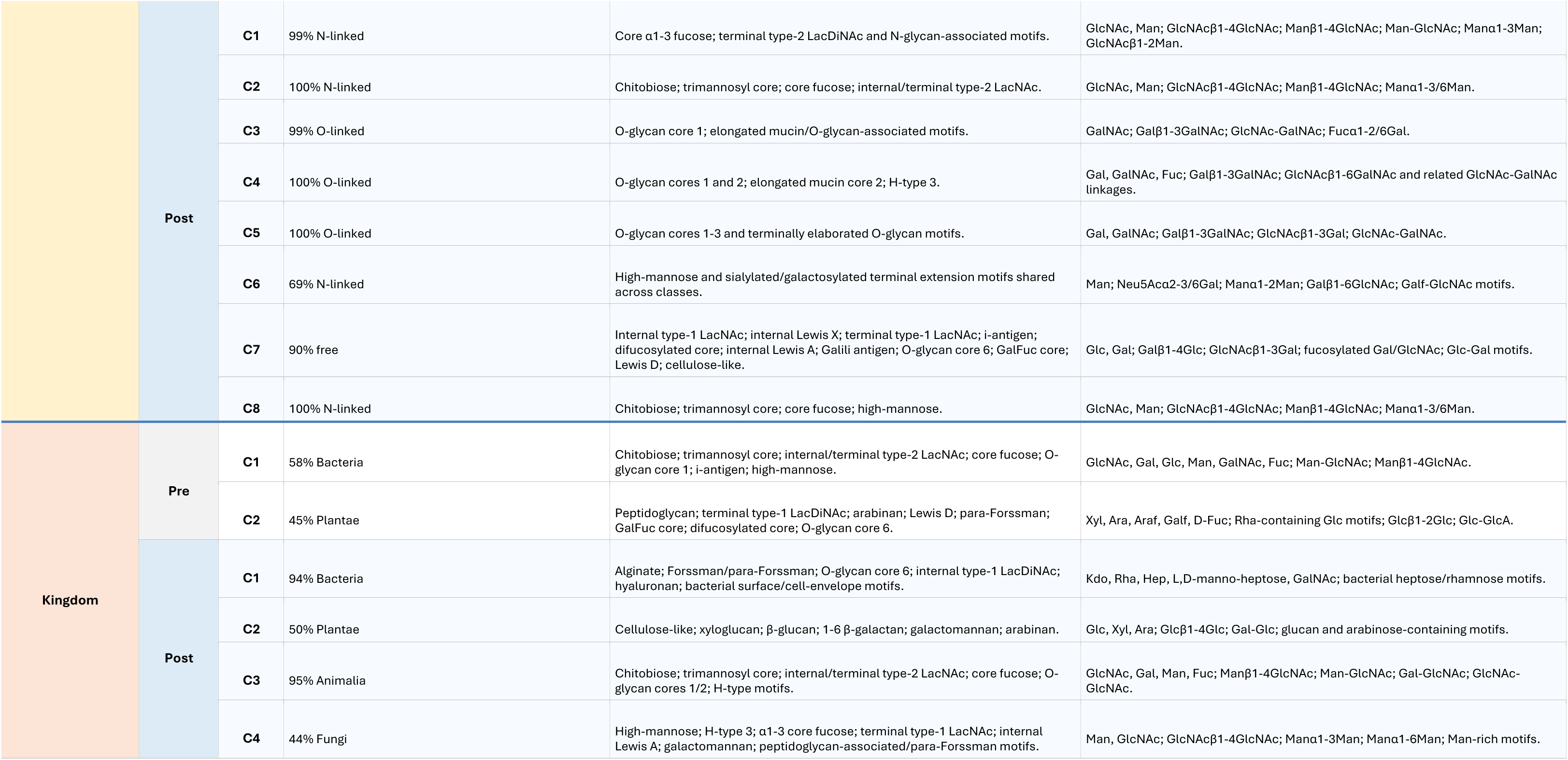
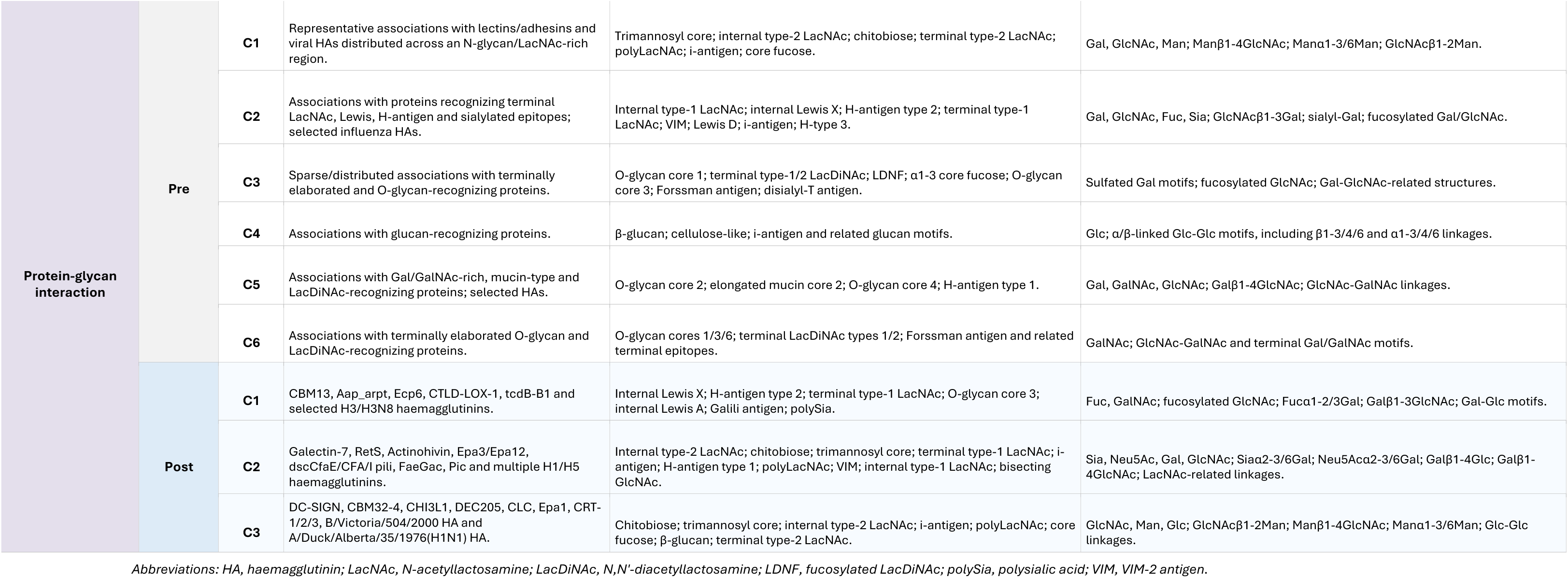
Cluster-level biological and structural enrichment before and after task-specific fine-tuning. Summary of the Leiden communities identified in the pretrained and fine-tuned ChemBERTa embedding spaces for immunogenicity, glycosylation-type, kingdom and protein-glycan interaction prediction. For each community, the table reports the dominant benchmark label or enriched protein associations, together with significantly enriched annotated glycan motifs, monosaccharides and glycosidic linkages. For the classification tasks, percentages indicate the proportion of glycans carrying the dominant label within each community. For protein-glycan interaction prediction, listed proteins showed enriched interaction scores for glycans assigned to the indicated community relative to all remaining glycans. Glycan, monosaccharide and linkage enrichments were calculated using cluster-versus-rest comparisons with multiple-testing correction and indicate relative over-representation rather than exclusive occurrence within a community

**Supplementary Table 6.** Experimental settings used throughout the GlycanMLBenchmark evaluation pipeline. Summary of the preprocessing, embedding extraction, downstream modelling and fine-tuning settings used throughout this study. The table reports the configuration employed for GlyLES conversion, foundation-model embedding extraction, protein embedding generation, Morgan fingerprint construction, downstream XGBoost and multilayer perceptron (MLP) predictors, task-specific fine-tuning, post-fine-tuning embedding extraction and benchmark aggregation. Unless otherwise stated, these settings were applied consistently across all benchmark tasks to ensure fair comparison between engineered features, frozen molecular representations, task-adapted chemical foundation models and specialist glycan-native models.

| Stage / model family | Benchmark scope | Hyperparameters and settings |
| --- | --- | --- |
| Preprocessing / GlyLES conversion | All benchmark datasets | GlyLES conversion to SMILES; deterministic processing; split columns preserved. No tuned model hyperparameters. |
| Static SMILES/sequence embeddings | ChemBERTa-zinc, ChemBERTa-77M, MoLFormer-XL, MoLFormer-c3 and MAMMAL | Batch size 256; model tokenizer context lengths as in Supplementary Table 3; mean and/or CLS pooling as configured; HF extraction uses model max-length padding/truncation; configured MoLFormer no-trunc variant retained. |
| Uni-Mol2 embeddings | Uni-Mol2 570M and 84M; mean and CLS representations | Batch size 32; max_atoms=128; hydrogens removed |
| Protein embeddings | Protein-glycan interaction benchmark | ESM2 facebook/esm2_t33_650M_UR50D; batch size 8; fixed protein embeddings combined with glycan representations for interaction prediction. |
| Fingerprint features | All four benchmark tasks; interaction also uses protein embeddings | RDKit Morgan Count and Morgan Bit fingerprints; radii 2-3; chiral and achiral variants; fpSize=2048. |
| Static XGBoost downstream models | Taxonomy, immunogenicity, glycosylation type and protein-glycan interaction | Binary: objective=binary:logistic, eval_metric=logloss, tree_method=hist. Multiclass: objective=multi:softprob with num_class set per task. Regression: objective=reg:squarederror, tree_method=hist. n_jobs=10; random_state=seed; |
| Static MLP downstream head | Static embeddings/fingerprints for all benchmark tasks | One hidden layer of 512 units with ReLU and dropout 0.1; AdamW; batch size 256; max_epochs 100; early stopping on val/loss. Learning rate: 1e-3 for taxonomy and glycosylation type; 1e-4 for immunogenicity and interaction. Patience: 10 for multiclass/regression and 5 for immunogenicity. |
| Finetuned Mods and Settings | ChemBERTa-zinc, ChemBERTa-77M, MoLFormer-XL and MoLFormer-c3 | AdamW lr=1e-4, weight_decay=1e-4; classifier/regression hidden_dim=512, dropout=0.1; freeze_lm/freeze_mol_encoder=False; precision=32; gradient_clip_val=1.0; accumulate_grad_batches=1; ReduceLROnPlateau factor=0.5, patience=2. Batch size 8 for classification tasks and 32 for interaction. |
| Post-fine-tuning embedding extraction | Fine-tuned backbones used as embedding generators | Batch size 128; CPU extraction; mean, CLS or last-token pooling according to saved run/config; MoLFormer outputs split by runner; no-trunc variants retained where configured. |
| Aggregation and reporting | Benchmark tables and figures | Metrics: macro-F1 for taxonomy/glycosylation type; AUPRC, F1 and accuracy for immunogenicity; Spearman, Pearson and MSE for interaction. Taxonomy summaries average over the eight taxonomic levels where used. |

